# Multiple roles of the polycistronic gene *tarsaless/mille-pattes/polished-rice* during embryogenesis of the kissing bug *Rhodnius prolixus*

**DOI:** 10.1101/667022

**Authors:** Vitória Tobias-Santos, Diego Guerra-Almeida, Flavia Mury, Lupis Ribeiro, Mateus Berni, Helena Araujo, Carlos Logullo, Natália Martins Feitosa, Jackson de Souza-Menezes, Evenilton Pessoa Costa, Rodrigo Nunes-da-Fonseca

**Affiliations:** Instituto de Biodiversidade e Sustentabilidade (NUPEM), Universidade Federal do Rio de Janeiro, Campus Macaé, Av. São José do Barreto 764, ZIP CODE: 27965-550; Institut de Génomique Fonctionelle de Lyon - CNRS UMR 5242, Lyon, France (32-34 avenue Tony Garnier 69007 Lyon); Laboratório de Biologia Molecular do Desenvolvimento, Instituto de Ciências Biomédicas, Universidade Federal do Rio de Janeiro, CCS Bl. F sala F2-031 Cidade Universitária Ilha do Fundão, ZIP CODE: 21941-970; Instituto Nacional de Ciência e Tecnologia em Entomologia Molecular - INCT-EM. INCT-EM, Rio de Janeiro, Brazil; Laboratório de Química e Função de Proteínas e Peptídeos, Centro de Biociências e Biotecnologia, Universidade Estadual do Norte Fluminense Darcy Ribeiro

**Author notes:** Corresponding author: RNdF.

## Abstract

Genes encoding small open-reading frames (smORFs) have been characterized as essential players of developmental processes. The smORF *tarsaless/mille-pattes/polished-rice* has been thoroughly investigated in holometabolous insects, such as the fruit fly *Drosophila melanogaster* and the red flour beetle *Tribolium castaneum*, while its function in hemimetabolous insects remains unknown. Thus, we analyzed the function of the *tal/pri/mlpt* ortholog in a hemimetabolous insect, the kissing bug *Rhodnius prolixus (Rp)*. First, sequence analysis shows that *Rp-tal/pri/mlpt* polycistronic mRNA encodes two small peptides (11 to 14 amino acids) containing a LDPTG motif. Interestingly, a new hemipteran-specific conserved peptide of approximately 80 amino acids was also identified by *in silico* analysis. *In silico* docking analysis supports the high-affinity binding of the small LDPTG peptides to the transcription factor Shavenbaby. *Rp-tal/pri/mlpt in situ* hybridization and knockdown via RNA interference showed a conserved role of *Rp-tal/pri/mlpt* during embryogenesis, with a major role in the regulation of thoracic versus abdominal segmentation, leg development and head formation. Altogether, our study shows that *tal/pri/mlpt* segmentation role is conserved in the common ancestor of Paraneoptera and suggests that polycistronic genes might generate order specific smORFs.

## Introduction

A large number of essential genes required for biological processes have been discovered by genetic screenings in model organisms such as the fruit fly *Drosophila melanogaster* (e.g.[1]). While most loci important for developmental processes were identified in these original screenings, recent genetic and expression analyses in *D. melanogaster* and in the beetle *Tribolium castaneum* showed that genes previously classified as putative noncoding RNAs encode functional small open reading frames (smORFs) or sORFs [2; 3]. smORFs, ORFs smaller than 100 amino acids, have been described as fundamental for several developmental processes of insects [4; 5; 6]. Although functional smORFs have been originally described in yeasts [7], gene prediction methods have, in general, discarded smORFs in genome-wide predictions (reviewed in [8]). Comparative genomic analysis of Drosophilid species showed an unexpected conservation of smORF containing genes, suggesting important biological roles for this new class of genes [9].

Later on, experimental analysis of conserved smORFs such as *sarcolamban (scl)*, a conserved peptide involved in Ca^2+^ uptake at the sarco-endoplasmic reticulum [10], and hemotin [11], a conserved phagocytosis regulator, provided further evidence that genes containing smORFs might constitute a reservoir of important players in metazoan genomes. In the past years, direct evidence of large-scale smORF translation has been obtained by ribosomal profiling of polysomal fractions in *Drosophila*, using the Poly Ribo-Seq technique [12]. This study was able to classify smORFs in two groups: “longer” smORFs of around 80 amino acids resembling canonical proteins, mostly containing transmembrane motifs, and shorter (“dwarf”) smORFs. These ‘dwarf’ smORFs are in general shorter (around 20 amino-acid long), less conserved and mostly found in 5’-UTRs and non-coding RNAs.

While bioinformatic studies point to hundreds or thousands of genes containing putative smORFs, only a few functional studies have been performed. In insects, the function of the smORF founding member *mlpt* was only investigated in detail in holometabolous insects such as the *D. melanogaster* and *T. castaneum* [4; 5; 6] and more recently in basally branching Diptera [13]and Lepidoptera [14]. In *T. castaneum mlpt* acts as a gap gene during the process of embryonic segmentation, regulating Hox genes and thoracic versus abdominal specification; knockdown for *mlpt* leads to embryos with multiple legs, *mille-pattes* in French [6]. In *D. melanogaster tal* was identified in a spontaneous mutant with defects in the distal part of the legs, the tarsus [5]. Independently, in the same year, Pri peptides were shown to be required for the F-actin organization during trichome morphogenesis; embryos lacking *pri* show external cuticle defects, resembling *polished rice* [4]. More recently, Mlpt peptides were shown to be regulated by ecdysone, defining the onset of epidermal trichome development, through post-translational control of the Shavenbaby (Svb) transcription factor [15]. Altogether, Mlpt peptides display context-specific roles and interactions within different developmental processes.

While in *D. melanogaster mlpt* early embryonic gene expression is segmental displaying a pair-rule pattern, mutants do not display segmentation or homeotic alterations as reported in *T. castaneum*. *D. melanogaster* embryonic mutant phenotypes include broken trachea, loss of cephalopharyngeal skeleton, abnormal posterior spiracles and lack of denticle belts, structures of late embryonic *mlpt* expression [5].

To clarify whether the segmentation function of *mlpt* is ancestral, but has been lost in the lineage giving rise to *D. melanogaster*, or whether it is a recently arisen specialization of *T. castaneum*, it is important to study *mlpt* function in other insect groups. Since hemipterans, as hemimetabolous, comprise the sister group of holometabolous insects (reviewed in [16]), a functional characterization of *mlpt* in the emergent hemiptera model, the kissing bug *Rhodnius prolixus* would be important [17]*. R. prolixus* available genomic and transcriptomic resources [18; 19; 20; 21; 22], an established embryonic staging system [23] and the recent availability of *in situ* hybridization and RNA interference techniques are great advantages of this model system [17].

Here we report interesting conserved and new functional aspects of the prototypic smORF *mlpt* gene. First, sequence analysis identified a new hemiptera-specific peptide in the polycistronic mRNA of *mlpt*, a gene conserved throughout the Pancrustacean clade. Second, molecular docking analysis indicates that the small peptide containing a LDPTG(L/Q/T)Y consensus motif interacts with the N-terminus of the transcription factor Svb as in *D. melanogaster*. Third, expression and functional analysis shows that the ortholog of *mlpt* acts during embryonic segmentation, regulating the transition between thoracic and abdominal identity, similarly to its role previously described in *T. castaneum.* Fourth, a conserved role in tarsal patterning was also observed. Overall, segmentation and tarsal patterning are conserved roles of *mlpt* and provide evidence that generation of new peptides from smORFs might constitute an underestimated mechanism for the evolution of new genes.

## Results

The recent description of several biologically important genes encoding smORFs among metazoans opened new avenues for molecular biology and functional genomics research. *mlpt* function has been largely investigated in holometabolous insects, while its function in hemimetabolous insects has not been fully investigated. In the current manuscript we describe a bioinformatic and functional analysis of *mlpt* function in the kissing bug *R. prolixus*.

### *mlpt* polycistronic gene and peptide distribution among arthropods

Previous BLAST searches for genes encoding *mlpt* peptides against different arthropod genomes and transcriptomes provided evidence that this gene containing smORFs is restricted to insects and crustaceans [5]. The increase in arthropod genome sequences in the past years e.g. [24; 25; 26] allowed us to perform a complete search in the available arthropod genomes. Using a non-stringent BLAST approach (see methods for details) we identified *mlpt* orthologs in available insect genomes (Sup. Table 1). These orthologs encode two to several copies of a peptide of 11-32 amino acids, containing a LDPTG(L/Q/T)Y consensus motif (Figure 1 A - red boxes, Figure 1B). These peptides have been previously shown to mediate the switch of the *svb* transcription factor from a repressor to an activator [27] via the ubiquitin-conjugating complex, UbcD6-Ubr3, and proteasome recruitment [28]. Remarkably, hemiptera *mlpt* orthologs not only contain two smORFs encoding the LDPTG(L/Q/T)Y consensus motif (Fig 1A-red boxes), but also a larger hemiptera-specific smORF of about 80 amino acids (Figure 1 A-green boxes and Figure 1 C). This peptide appears to be smaller and less conserved in basally branching hemipteran species with circa 60 amino acids in the genome of the pea aphid *Acyrthosiphon pisum* and *Pseudococcus longispinus* and larger than 80 amino acids in triatomes such as *Rhodnius prolixus* and *Triatoma pallidipennis.* This new predicted peptide is potentially translated from a small CDS generated through post-transcriptional regulation, since it is interspersed by two introns in its respective gene in at least three hemipteran species. (Sup. Fig. 1). We named this new peptide as smHemiptera due to its restricted phylogenetic distribution. Interestingly, a polycistronic transcript containing all the aforementioned smORFs was identified in a digestive tract *R. prolixus* transcriptome ([21] and our own observations).

**Figure 1:**
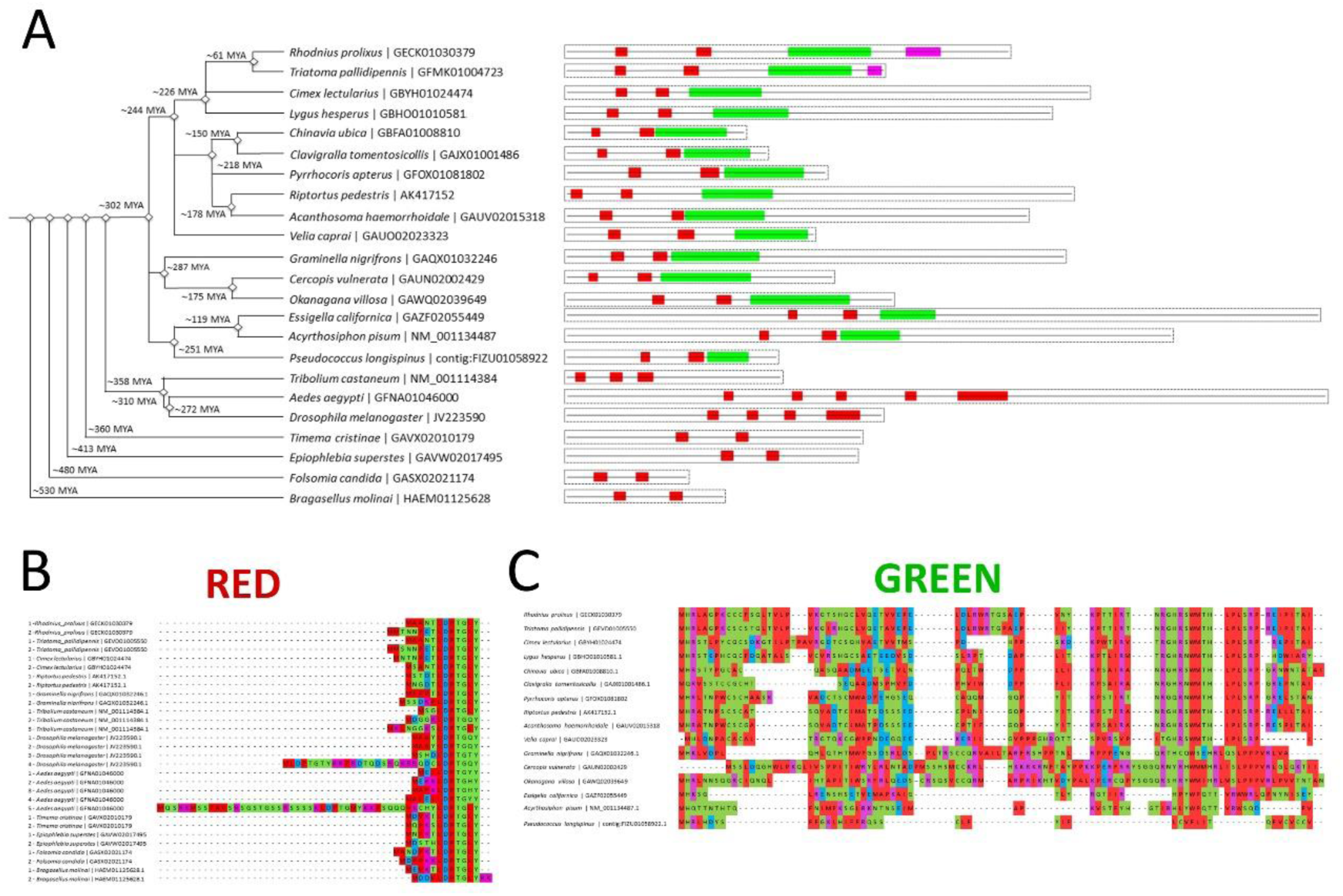
*mlpt* evolutionary analysis and putative regulation. A) Left - Phylogenetic representation of *mlpt* distribution among arthropod species and approximately time of node speciation. the online server TimeTree [75]. *mlpt* polycistronic mRNA organization. White boxes comprise the whole polycistronic mRNA containing the smORFs encoding peptides displayed in red, green and pink. B) Alignment of the previously described small peptide (red box in Figure 1A) containing an LDPTG domain [4; 5; 6], conserved in several arthropod species and whose number of paralogs in a single polycistronic gene varies among arthropod species. C) Alignment of a hemipteran specific peptide of about 80 amino acids (green box in Figure 1A. The pink smORF encoded peptide is only observed in triatomines and its alignment was omitted for simplicity.

### Interaction mode prediction, hotspots and binding affinities of Shavenbaby-Mlpt complexes

Previous work in *D. melanogaster* identified the 31 N-terminal residues of the transcription factor (TF) *svb* as a Mlpt-dependent degradation signal, or degron [28]. In order to investigate the interaction modes between the peptides encoded by the polycistronic mRNA of *mlpt* and the transcription factor Shaven baby (Svb) from *D. melanogaster* (Dmel-Svb) and *R. prolixus (*Rpro-Svb) 3D structures were predicted and docking assays performed. Secondary structure analysis indicates that both Dmel-Svb and Rprol-Svb display mainly typical features of intrinsically disordered proteins, as previously suggested [28]. The disordered regions appear in large regions of both proteins, being interspersed by small alpha-helices and some very short beta-sheets (Figure 2 A,B). Smaller peptides containing LDPTG(L/Q/T)Y motifs, Rprol-pptd1, Rprol-pptd2 and Dmel-pptd1 were considered disordered, as well as smHemiptera peptide (Figure 1C).

**Figure 2:**
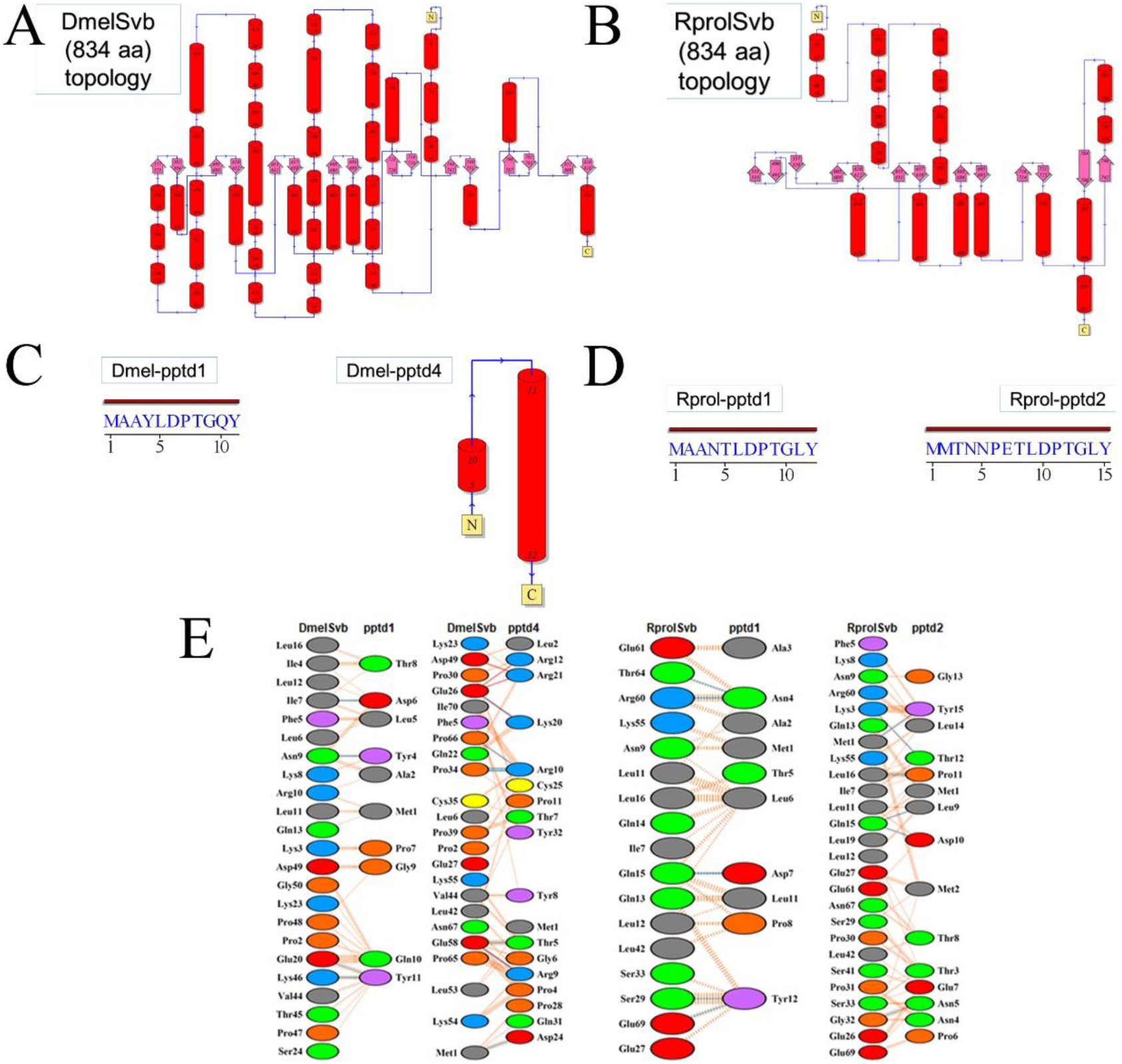
Docking and *in silico* mutation analysis of the interaction between Svb and Mlpt peptides. (A,B) *Drosophila melanogaster* (A) and *Rhodnius prolixus* (B) protein topology. (C,D) Predicted topology of Mlpt peptides in *D. melanogaster* (Dmel) and *R. prolixus* (Rprol). (E) Predicted interactions between Svb proteins and small peptides containing the LDPTG(L/Q)Y domain.

In contrast, Dmel-pptd4, a peptide with 32 aa, and previously described as non-translated in *Drosophila* cells [4; 5], showed two alpha-helices, intercalated by disordered regions (Figure 2 C,D). Docking assays between Svb and the small peptides containing LDPTG(L/Q/T)Y motifs identified the main interaction types at the interface between the complexes (Figure 2 E), as well as their interaction affinities (Sup. Table 2). According to the *in silico* data, the interaction between Dmel-pptd4 presented five salt bridges, seven hydrogen bonds and 127 non-bonded contacts. Its binding affinity parameters indicate a high affinity to the transcription factor Svb. The hotspots analysis, on amino acids located at the protein-protein complex interface and evaluated as the most critical for its maintenance, are listed on Supp. Table 2. The binding affinity change upon mutations indicate that, for the transcription factors (Dmel-Svb and Rprol-Svb), the destabilization of complexes tends to occur mainly through double mutations Supp. Fig. 1-2). On the other hand, for the small peptides (Dmel-pptd1 and 4, Rprol-pptd1 and 2), single mutation effects can be enough to weaken the protein-peptide complex, this effect being significantly increased with double mutations. Interestingly, the most conserved amino acids Lysine (Lys), D (Asp) and Y(Tyr) residues of Dmel-pptd1, Rprol-pptd1 and 2 (Figure 1 B) display multiple interaction residues with Svb in *D. melanogaster* and *R. prolixus* (Figure 2 E). *In silico* data did not detect high affinity interactions between smHemiptera and Rp-Svb N-terminal region, suggesting that this hemipteran specific peptide might show different molecular partners. Altogether, our molecular docking results corroborate previous experimental data [28] and suggest that similar amino acid residues in Svb and Mlpt are essential in both species for protein-peptide complex formation.

### Rp-mlpt regulatory region shows putative binding sites for Ecdysone Receptor

Recently, *mlpt* expression was shown to be directly activated by the ecdysone pathway via direct binding of the nuclear receptor EcR to cis-regulatory sequences of *mlpt* [15]. We searched upstream, intronic and downstream sequences of *D. melanogaster mlpt (Dm-mlpt)*, *T. castaneum mlpt (Tc-mlpt)* and *R. prolixus mlpt (Rp-mlpt)* for the occurrence of *D. melanogaster* Ecdysone Receptor binding sites (Figure 3). As expected, one EcR binding site is observed in the regulatory region of *Dm-mlpt*; this region was previously shown to be directly bound by Ecdysone in ChIP-seq analysis of *D. melanogaster* [15]. Searches in the *T. castaneum mlpt* locus with the same TF binding site showed four putative binding sites for EcR, upstream of the transcription start site (TSS) and one binding site several kilobases downstream, suggesting that a regulatory region responsive to ecdysone might reside upstream the TSS in this beetle. While the loci of *D. melanogaster* and *T. castaneum* display similar genomic size, *R. prolixus mlpt* locus is much larger due to the large intronic regions which separate the three distinct coding regions (CDS) of the new hemipteran-specific peptide smHemiptera. Analysis of this locus in *R. prolixus* showed that six EcR binding sites occur particularly close to smHemiptera. This result suggests that ecdysone might also regulate *mlpt* expression during *R. prolixus* developmental transitions.

**Figure 3:**
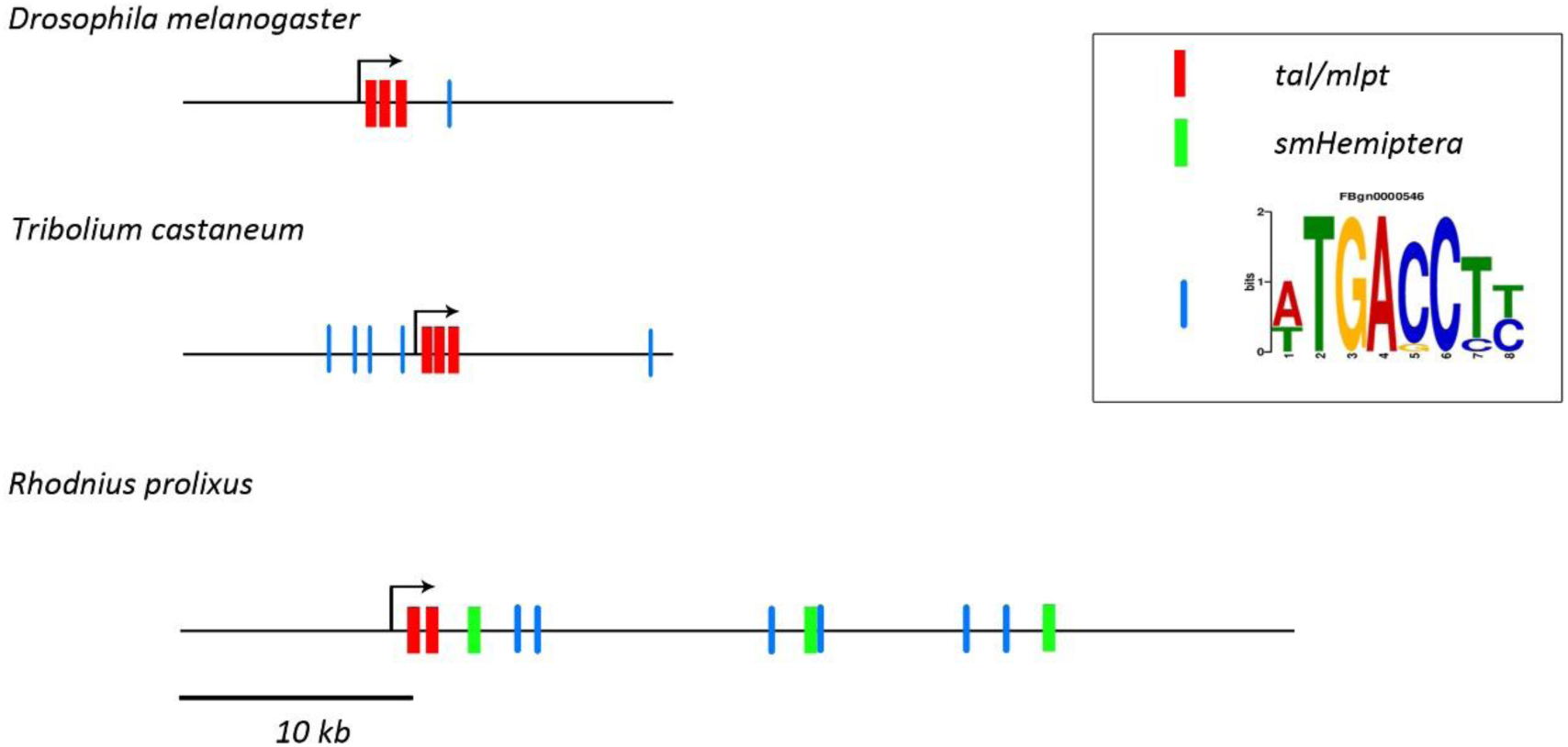
Regulatory landscape of *mlpt* in *D. melanogaster*, in *T. castaneum*, and in *R. prolixus*. Genomic loci of the gene *mlpt* from *D. melanogaster, T. castaneum* and *R. prolixus*. Red boxes represent ORFs containing LDPTG(L/Q)Y domains, green boxes represents the polycistronic Hemiptera peptide (smHemiptera). The putative EcR binding sites (FBGn0000546) are highlighted in blue.

### Rp-mlpt spatial expression pattern and relative expression during Rhodnius prolixus embryogenesis

*mlpt* embryonic expression was originally described in the beetle *T. castaneum* and has been characterized by a very dynamic zygotic patterning [6]. In *D. melanogaster*, *mlpt* expression was correlated with tissue folding, acting as a connection between patterning and morphogenesis [5]. Early expression of *D. melanogaster mlpt* starts as seven blastodermal stripes and a cluster of cells in the anterior part of the embryo, although segmentation is not affected in *mlpt* mutants. Later, after segmentation, *mlpt* is present in the trachea, posterior spiracles, pharynx, hindgut, and presumptive denticle belts. *D. melanogaster mlpt* mutants display reduced cuticular structures and ectopic expression of *mlpt* in the head induces extra skeleton components [5].

*R. prolixus mlpt* distinct expression can be observed at stage 1B, between 6-12 hours after egg laying (AEL), in an anterior domain (Figure 4 A), which corresponds to the presumptive head lobes, followed by the appearance of a posterior expression domain shortly before the posterior invagination at stage 2A (Figure 4 B-12-18 hours AEL). Hemipterans, such as *R. prolixus*, undergo an embryonic inversion movement called anatrepsis [29]. At stage 3A (24-30 hours AEL), during this inversion movement (Figure 4 C,C’), and at the beginning of germ band elongation at stage 3B (Figure 4 D,D’-30-36 hours AEL), *Rp-mlpt* is expressed on the newly formed thoracic and abdominal segments, but not at the posterior-most region, the segment addition zone (SAZ) [30]. At stage 4 during gem band elongation *Rp-mlpt* expression remains in front of the SAZ and in an anterior expression domain of the head (Figure 4 E-J −36-48 hours AEL). At stage 5 (48-60 hours AEL), shortly before the end of segmentation and when segmental buds are about to appear, *Rp-mlpt* is expressed only at the posterior-most region (Figure 4 K – 48-60 hours AEL). Between stage 5 and stage 6 (60-72 hours), abdominal segmentation is completed and *Rp-mlpt* expression is observed at the tips of the legs and in the head lobes (Figure 4 L). Between stage 7 (72-84 hours AEL) and stage 8 (84-96 hours AEL), which is characterized by leg proximal-distal axis growth and abdominal segments thickening, an expression at the lateral, future dorsal embryonic region can be visualized (Figure 4 M). At stage 9 (96-108 hours AEL), at the beginning of germ band retraction, *Rp-mlpt* is only observed at the antenna, an organ undergoing extensive increase in size during stage 9 and 10 (Figure 4 N, [23]). Altogether, *mlpt* expression is highly dynamic, suggesting multiple roles during *R. prolixus* embryonic development.

**Figure 4:**
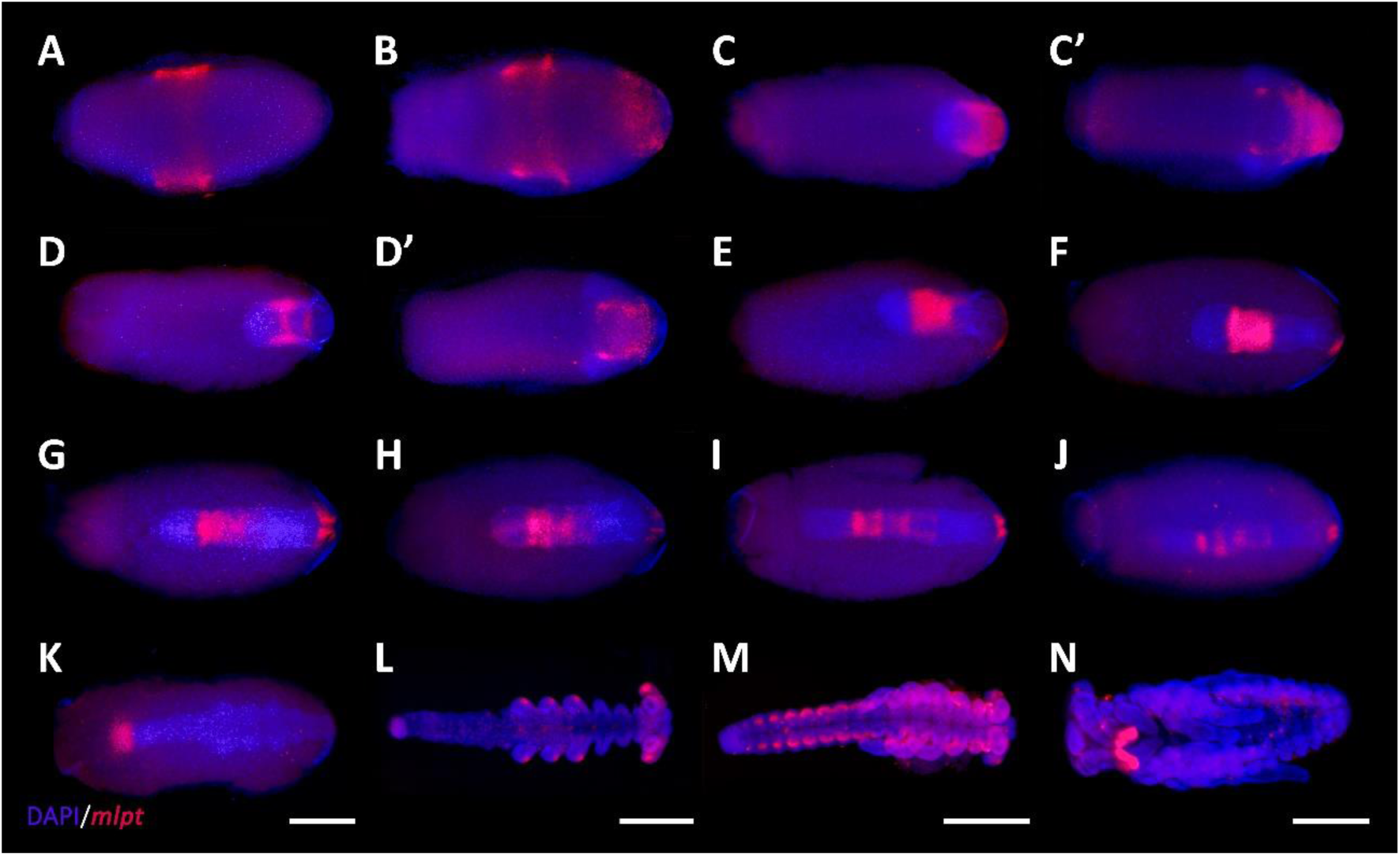
*mlpt* expression during *R. prolixus* embryogenesis. Anterior of the egg to the left in all panels. During the processes of anatrepis and katatrepsis embryo orientation shifts along the AP and DV axis (see [17] and [23] for details). *mlpt* expression is displayed in falsecolored red and DAPI in blue. Ventral egg view (A,B,C’,D’,G,H,I). Eggs dissected from the yolk (L,M,N) (A) Stage 1B, approximately 10 hours after egg laying (AEL) *mlpt* expression is detected as a broad middle-anterior domain, which overlaps with to the future head. (B) A few hours later (stage 2A-12-18 hours AEL) besides the anterior domain, a posterior domain is also observed. (C,C’,D,D’) At stage 3A (24-30 hours AEL) during anatrepsis, the posterior expression domain is at the surface (ectoderm) but does not extend towards the posterior region of the germ band. (E-J) At stage 3B (30-36 hours AEL) the expression remains anterior to the posterior region and in a small bilateral domain in the head and starts to fade as elongation is finished. (K) Stage 4 (36-48 hours) At beginning of head and thoracic segmentation, expression is observed at the posterior tip. (L) Stage 5 (48-60 hours AEL) during leg growth a staining is specifically observed at the tips of the legs and in the head lobes. (M) At stage 7 (72– 84 hours AEL) abdominal segmentation is evident and expression is observed at lateral segmental cell clusters potentially corresponding to peripheral neurons as previously suggested for *T. castaneum* [6]. (N) Stage 9 (96–108 hours AEL) during germ band retraction expression is restricted to the antennae.

To investigate if levels of expression of *mlpt* also vary along embryonic development, we investigated its expression via Real-Time PCR. Small changes of *Rp-mlpt* expression were observed between non-fertilized (0-6 hours NF) and early fertilized eggs (0-6 hours AEL), while a ten-fold increase in expression was detected by the analysis of 36-48 hours AEL eggs (Figure 5 A). Comparatively, expression of the transcription factor *Rp-svb*, a known interaction partner of *mlpt* in other organisms[27] increased its expression only 40% upon fertilization and decreased 55% at 36-48 hours AEL, when compared to non-fertilized eggs (0-6 hours NF) (Figure 5 B).

**Figure 5:**
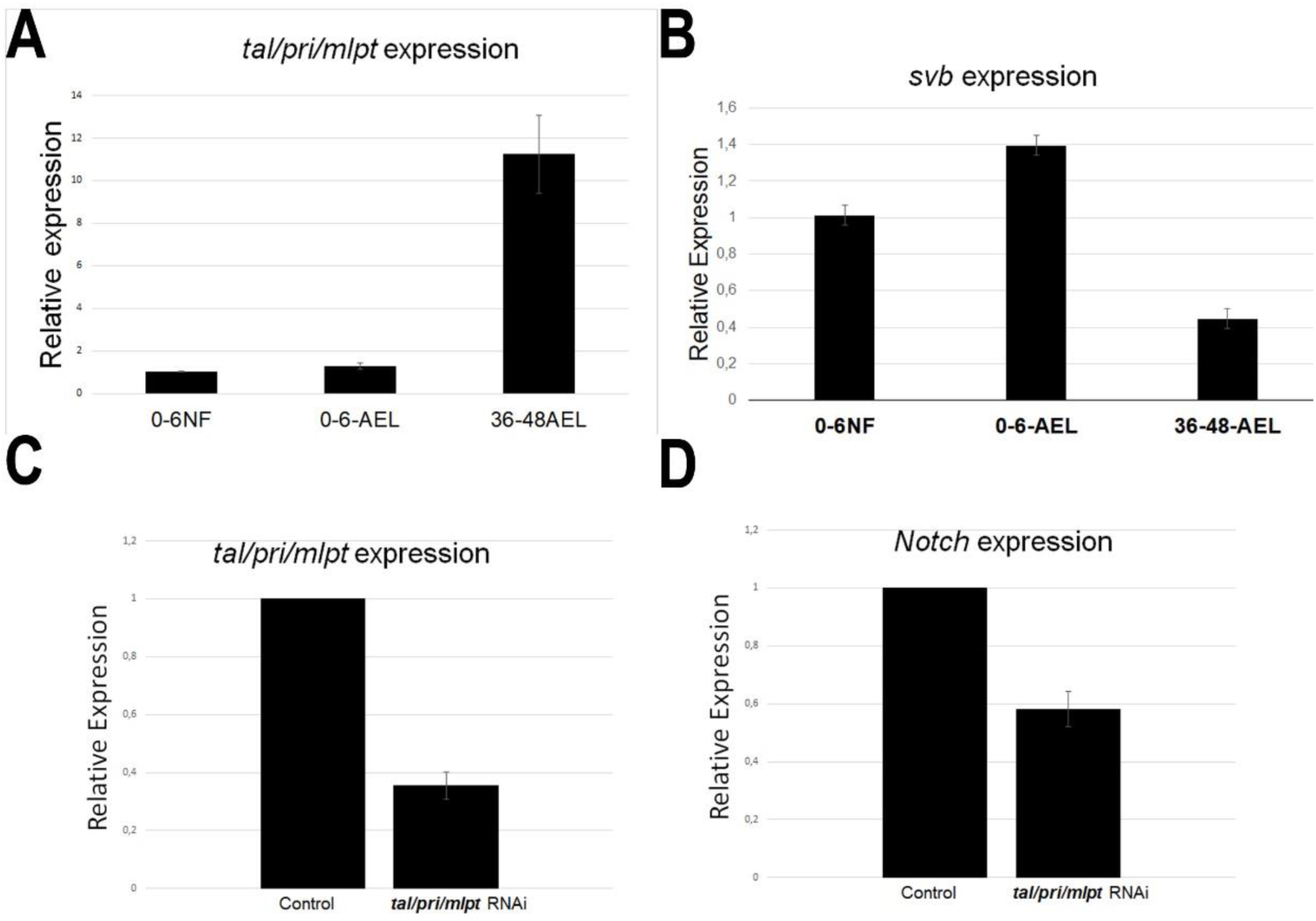
Temporal expression analysis and RNAi downregulation of *mlpt* during *R. prolixus* embryogenesis. (A,B) *mlpt* relative expression in non-fertilized (0-6NF), fertilized (0-6AEL) and 36-48 hours after egg lay (AEL). (C,D) Relative expression of *mlpt* (C) and Notch (D) in control and *mlpt* RNAi embryos.

### Rp-mlpt parental RNA interference is efficient and leads to a series of knockdown phenotypes

Since *Rp-mlpt* gene displays a complex expression patterning during embryogenesis, we sought to analyze its function by parental RNA interference (pRNAi) in *R. prolixus*, as previously described [23]. In this method, females are injected with double-stranded (dsRNA) synthesized against the gene of interest and phenotypic effects are evaluated in the offspring. Eggs collected from females injected with the control dsRNA (dsNeo) and the *Rp-mlpt* dsRNA showed a decrease up to 60-70% in *Rp-mlpt* expression in the latter dsRNA group, validating the knockdown (Figure 5 C). Increase in *Rp-mlpt* dsRNA injection concentration up to six micrograms per microliter (6µg/µl) did not further increase knockdown efficiency (data not shown). We also evaluated *Rp-Notch* expression, a receptor from a signaling pathway previously shown to interact with *mlpt* during leg formation [31], and embryonic segmentation in some insect species [32]. *Rp-Notch* expression decreased around 40% after *Rp-mlpt* in embryos after parental RNAi, suggesting a transcriptional regulation of *Rp-Notch* (Figure 5 D).

Upon egg collection and fixation, nuclear DAPI staining was performed and a plethora of embryonic mutant phenotypes were observed. A first phenotypic class of *Rp-mlpt* RNAi embryos comprises embryos with a wild-type number of segments, but with thoracic or gnathal segmental fusion. Embryos with this morphology were observed at stage 5 and 7 (Figure 6 B,B’, C,C’). A second phenotypic class was constituted by embryos with four pairs of legs in the *Rp-mlpt* RNAi instead of the three pairs observed in controls (Figure 6 E,E’). Last, a third phenotypic class of *Rp-mlpt* RNAi embryos shows improper germ band elongation and lack of the posterior region at stage 3 (Figure 6 A,A), which is more evident at stage 9, when lack of abdominal segmentation and fused legs were observed (Figure 6 D,D’).

**Figure 6:**
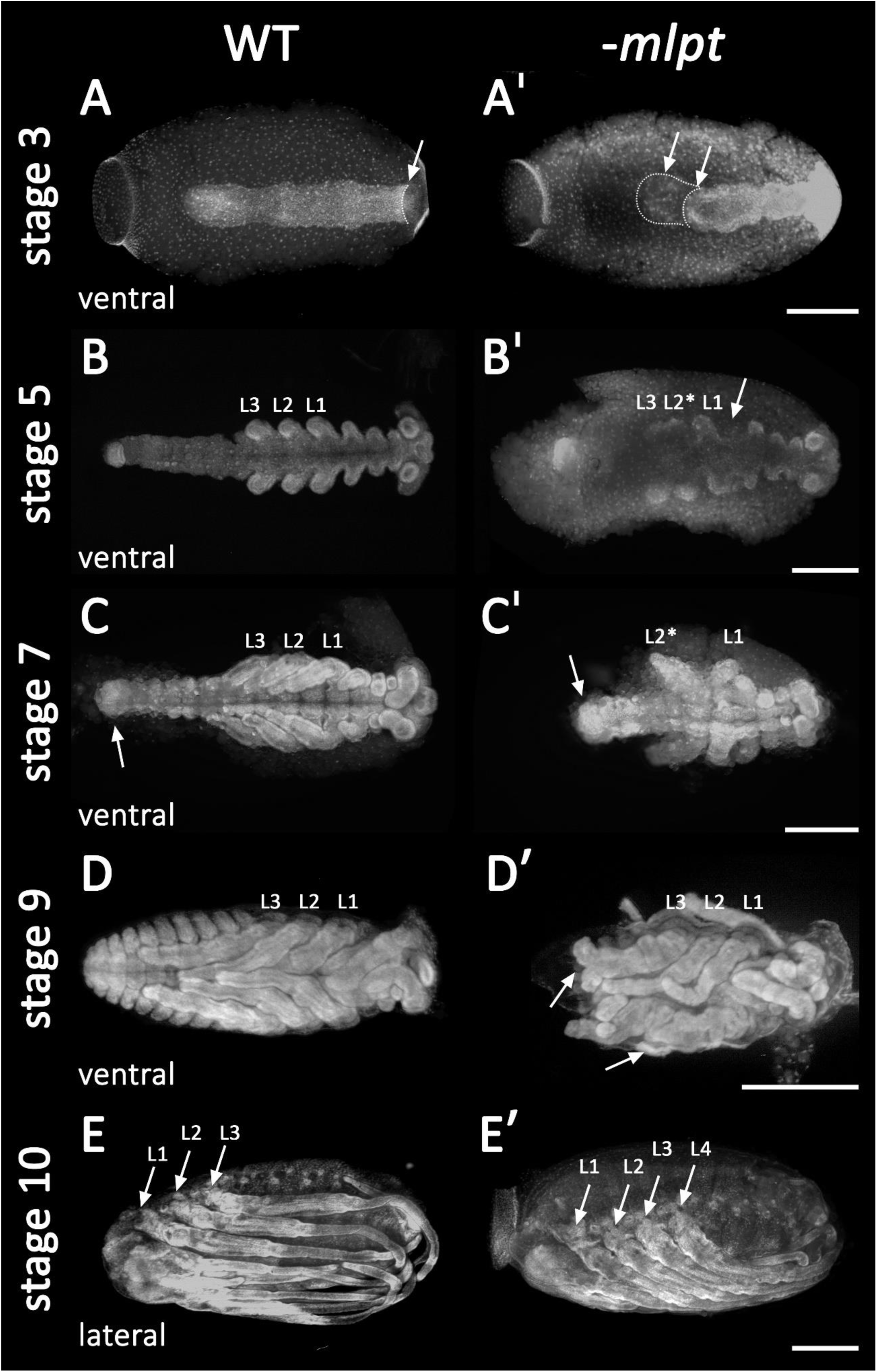
*mlpt* knockdown segmentation defects. DAPI staining of control and *mlpt* RNAi embryos. Anterior region of the egg towards to the left in all panels. (A,A’) Stage 3A. Germ band elongation appears irregular in the RNAi embryo when compared to the controls. (B,B’) Stage 5. Irregular thoracic and gnathal segmentation in the RNAi embryo. (C,C’) Stage 7. Thoracic segmental fusion and improper abdominal in the RNAi embryo segmentation when compared to the control. (D,D’) Stage 9. Third thoracic segment fused lack of abdominal segmentation and dorsal closure when compared to the control (E,E’) Stage 10. Late *mlpt* RNAi embryo showing four pairs of legs when compared to the three pairs of legs in the control. *R. prolixus* embryonic staging system was described in [23]

These phenotypic classes can also be observed by dissection of embryos at very late stages when nuclear DAPI staining is not possible due to cuticle formation (Figure 7 A-D). Thoracic red pigmentation can be used as a proxy to compare wild-type and RNAi embryos. In the first *Rp-mlpt* RNAi phenotypic class, embryos with two pairs of thoracic segments can be identified, indicating improper thoracic segmentation (Figure 7 B, B’, B’’, B’’’). In the second class, four thoracic segments are present (Figure 7 C, C’, C’’, C’’’), while in the third class, the strongest knockdown phenotype, the late embryo displays impairment of germ band elongation and most abdominal segmentation is lost, while dorsal closure does not take place (Figure 7 D, D’, D’’, D’’’).

**Figure 7:**
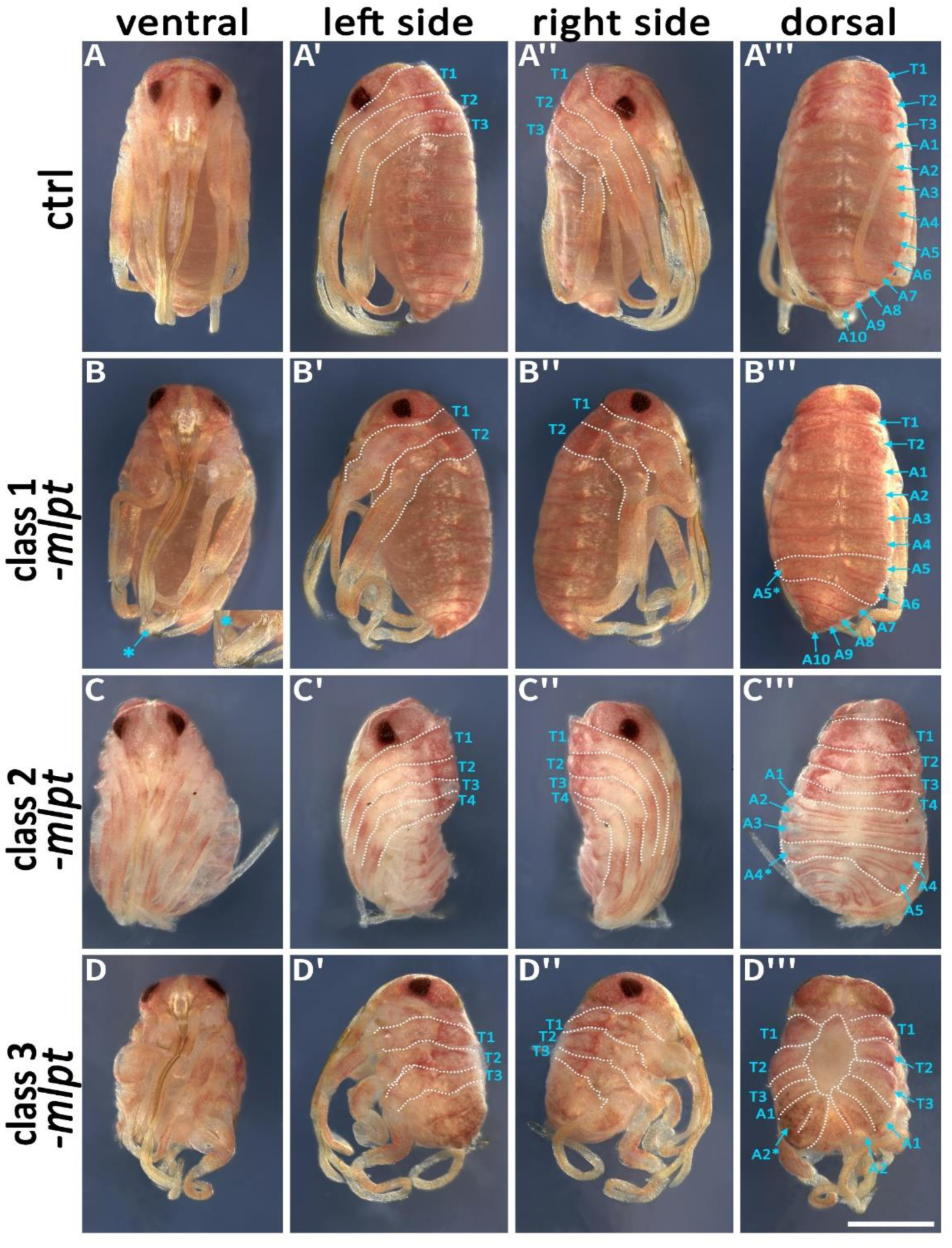
Phenotypic classes of *mlpt* RNAi late embryos. *mlpt* knockdown embryos were classified in three phenotypical classes when compared to the control. Class 1 - Embryos displaying only two thoracic segments and distal leg duplications (asterisk). Abdominal segmentation appears normal with the exception of some asymmetrical segmental divisions on left and right sides (asterisk) (20 out of 45 embryos). Class 2 - Embryos displaying four thoracic segments and irregular abdominal segmentation. (15 out of 45 embryos). Class 3 - Shorter embryos only displaying thoracic segments and fewer abdominal segments then controls (10 out of 45 embryos).

Interestingly, a few *Rp-mlpt* nymphs display unilateral segmental fusion of first and second thoracic segments leading to abnormal distal leg morphology (Figure 8 A-D). Further detailed analysis of nymphal legs showed abnormal morphologies in knockdown embryos (Figure 9). Distal regions of legs in wild-type *R. prolixus* are composed of a stereotypic pattern of tibia, tarsus and a pair of claws (Figure 9 A). *Rp-mlpt* RNAi legs show inappropriate tarsal formation (Figure 9 B), duplication of claws (Figure 9 C) and tarsus (Figure 9 D), and improper growth and lacking the distinction of tarsus and claws (Figure 9 E). Altogether, our results show that *Rp-mlpt* is important for the distinction between thoracic and abdominal segments, for the process of germ band elongation and distal leg patterning.

**Figure 8:**
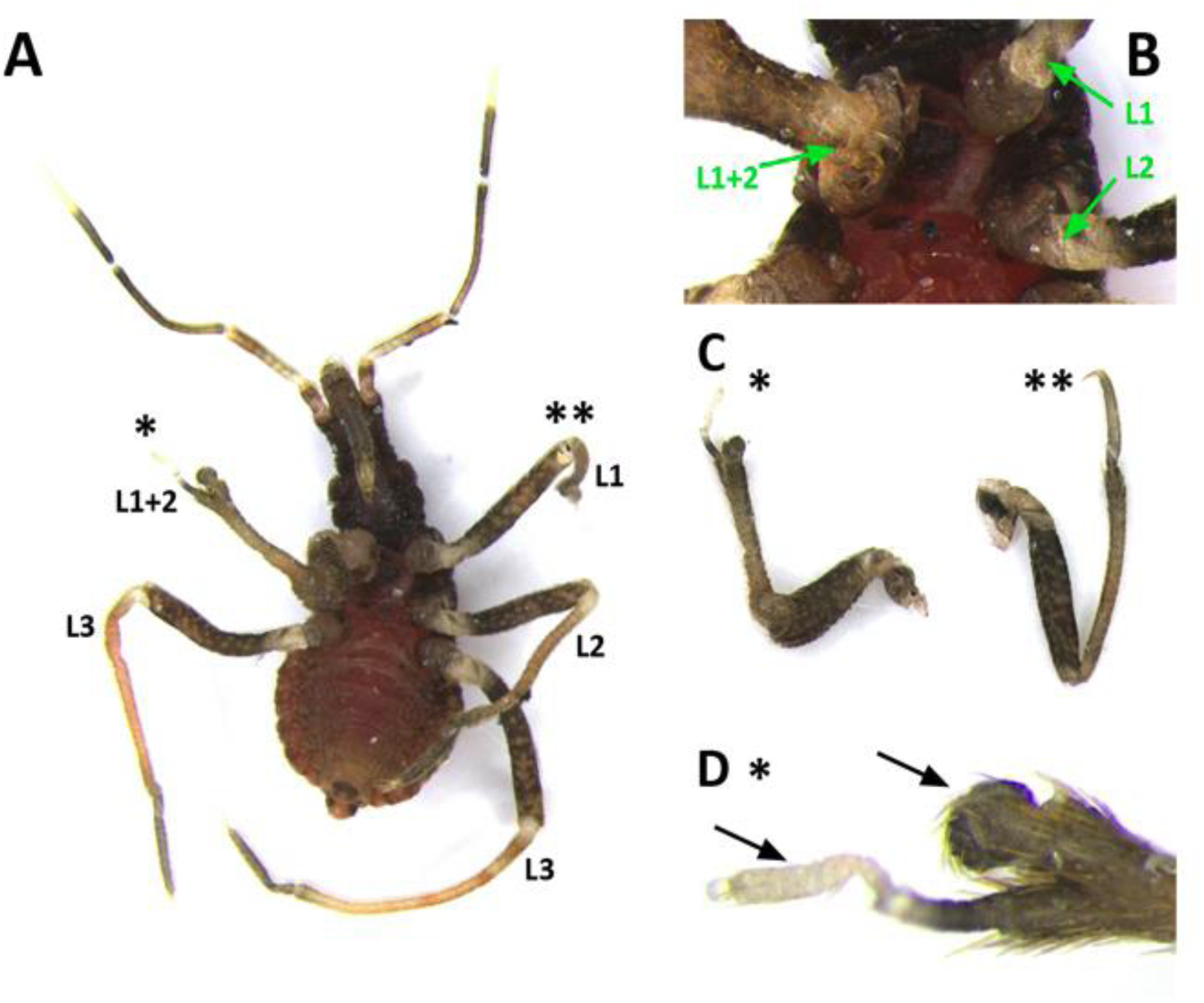
Segmental fusion after *mlpt* RNAi. (A-D) *mlpt* RNAi nymph showing unilateral leg fusion of L1 and L2. (B) Higher magnification of the proximal region of the segmental fusion. (C,D) Higher magnification of distal regions of fused legs show morphologies that deviate from the wild-type pattern. Tb - tibia, Ts - tarsus, C - claws.

**Figure 9:**
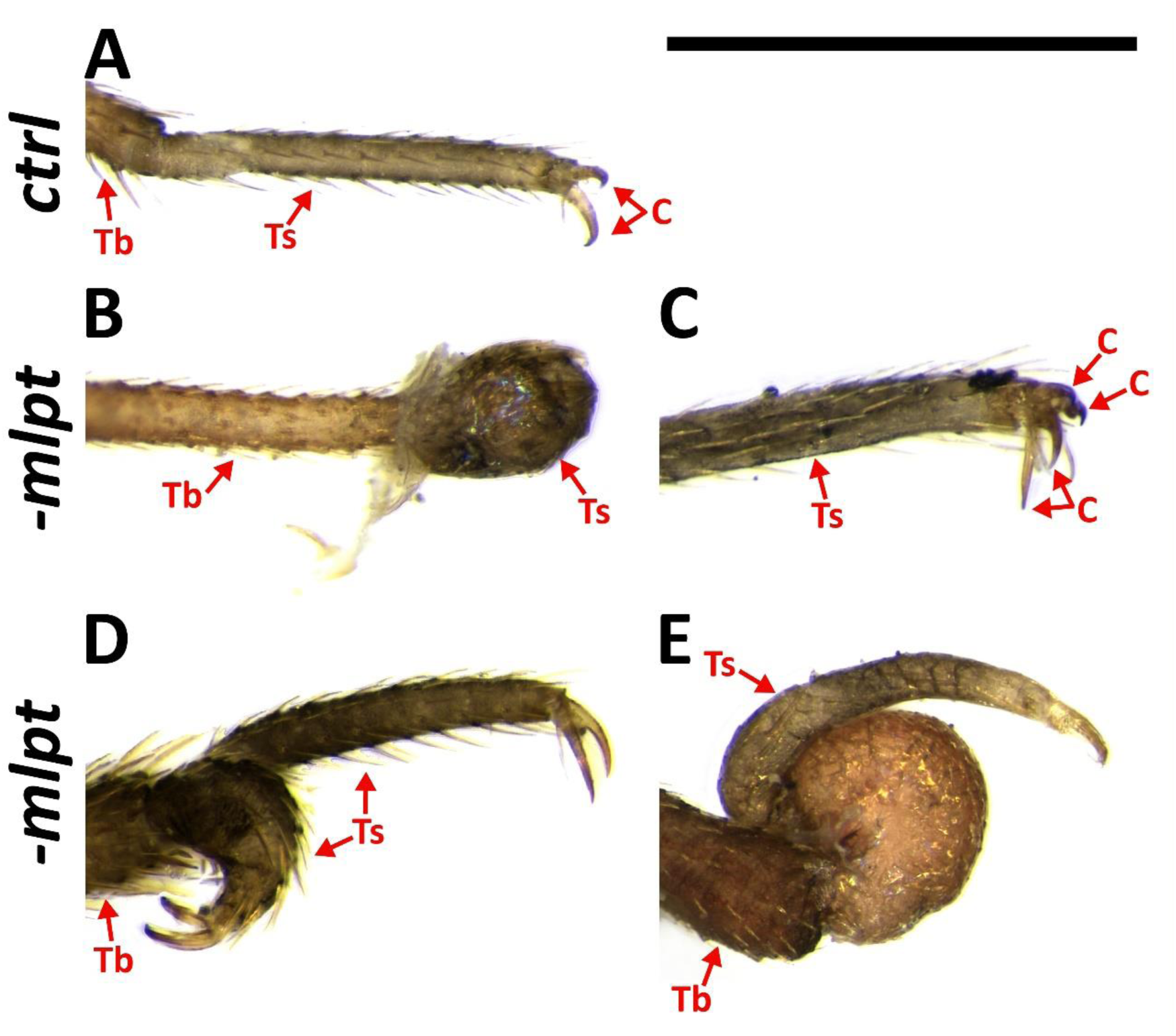
Legs are severely affected after *mlpt* RNAi. (A) Nymph control leg - Distal segments. (B-E) Phenotypic series of nymph *mlpt* RNAi legs. Several phenotypic changes from the wild-type pattern are observed after *mlpt* RNAi such as lack of distal regions such as tarsus (Ts) and claws as highlighted in B. (C,D) Duplications in claws (C) and tarsus (D) were observed.(E) Improper distal growth was also observed. Tb - tibia, Ts - tarsus, C - claws.

### Rp-mlpt RNAi leads to segmental fusion and impairment of head formation

Reduction in *Rp-mlpt* expression affected segment formation with clear phenotypic effects in the thorax and abdomen, before posterior invagination and at later segmentation stages (Figures 6 and 7). Since reduction or increase in the number of thoracic segments was observed in late embryonic stages after *Rp-mlpt* RNAi, we evaluated the expression of the ortholog of the segment polarity gene *hedgehog (hh)* in control and *Rp-mlpt* RNAi embryos. Like in other insects, the posterior region of every segment expresses *Rp-hh* during embryonic segmentation, and thoracic and abdominal segments can be identified by their differential widths (Figure 10). A phenotypic series of *Rp-mlpt* RNAi embryos shows that the distance between segmental stripes is reduced and that the distinction between thoracic and abdominal segments is less clear. Remarkably, in apparently stronger knockdown phenotypes, lack of the anterior-most segments of the head was also observed (Figure 10, asterisk). Altogether, *Rp-hh* expression analysis shows that *Rp-mlpt* is essential for proper thoracic and abdominal segmental distinction and for head formation.

**Figure 10:**
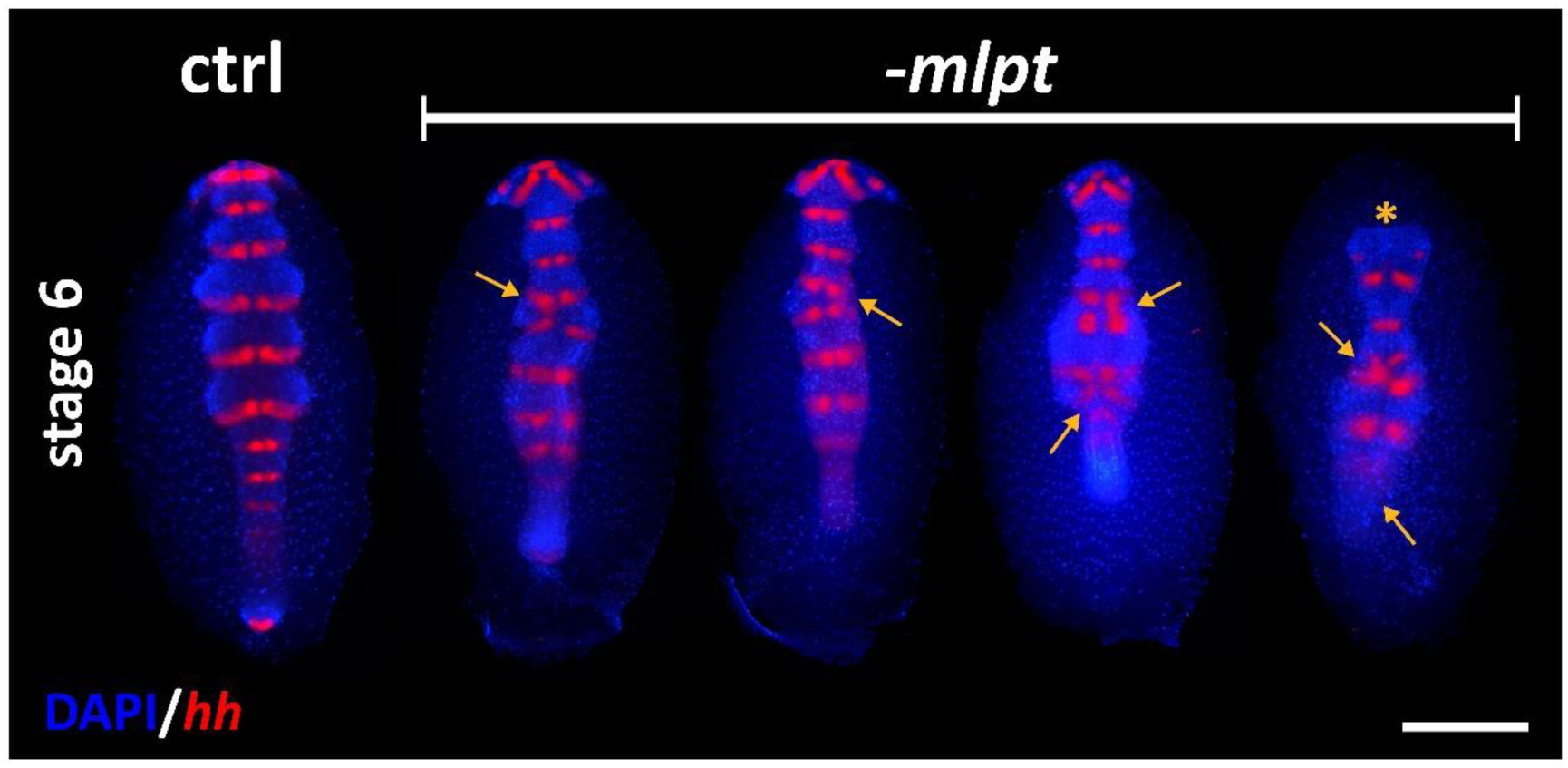
Control and phenotypic series of *mlpt* RNAi embryos. *in situ* hybridization for *hedgehog* (*hh*) and nuclear counterstaining with DAPI. Weakest phenotype (left) shows an apparent segmental fusion at the thoracic region. Progressive stronger phenotypes show fusion also in the abdominal segments. In the strongest phenotype (right most) the head is affected (asterisk) and germ band elongation did not occur. Arrows highlight segmental fusions.

### Comparison of knockdown phenotypes of mlpt and other developmental genes

Previous functional analysis of *mlpt* in the *T. castaneum* and *D. melanogaster* showed that this gene is required for early patterning in the beetle but not in the fruit flies. In beetles *mlpt* acts as an gap gene, while its role in fruit flies is restricted to later (post-segmental) embryonic stages [4; 5; 6; 27; 33].

To address if *mlpt* knockdown is comparable to the loss of other developmental genes, we performed parental RNAi against the orthologs of the gap genes *Krüppel (Kr)*, and *giant* (*gt*). Expression of the Hox gene *proboscipedia* (*pb*) in the third gnathal segment is comparable in control and *mlpt* RNAi embryos (Figure 11). In contrast, *pb* expression is duplicated upon *Rp-kr* RNAi and largely diminished upon *Rp-gt* RNAi (Figure 11). Analysis of the RNAi phenotype of the *mlpt* target, the transcription factor *svb* [27; 33], did not show any alteration on the expression of *pb* either, although germ band elongation was clearly affected (Figure 11).

**Figure 11:**
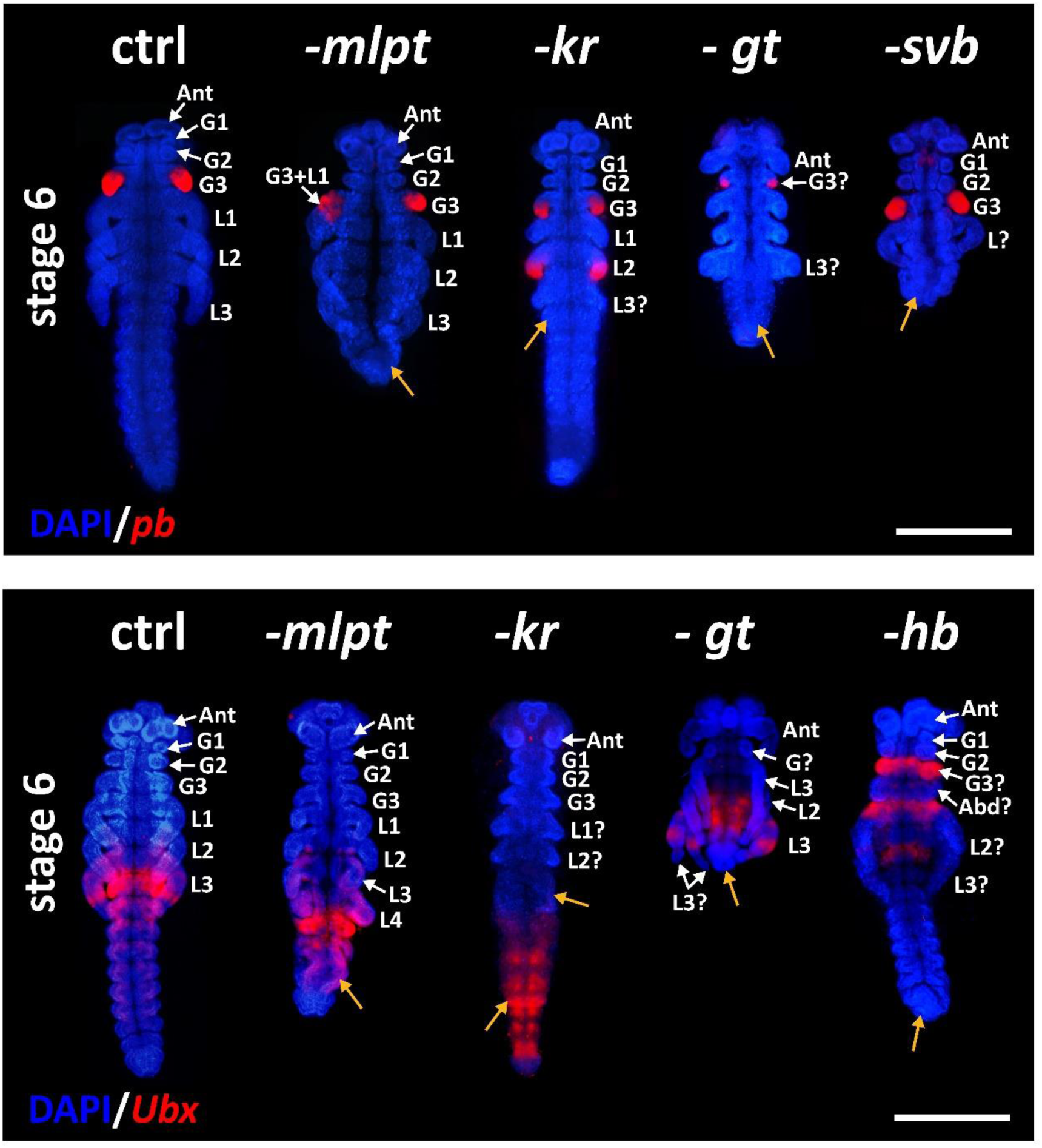
Hox gene expression in knockdown embryos of *mlpt* and other developmental genes. Upper row - In control embryos at stage 6, *pb* is expressed in the third gnathal segment, corresponding to the prospective labium. After *mlpt* RNAi *pb* is either non-affected (right side) or an irregular shape (lef side). Duplications or lack of staining were not observed after *mlpt* RNAi. *Kr* RNAi embryos show duplication patterns, *gt* RNAi embryos show a smaller group of expressing cells and the lack of anterior gnathal segments. *svb* RNAi displays clear segmentation deviations from the wild-type pattern, but does not display changes in *pb* staining. Lower row - *Ubx* expression in control embryos. Expression is stronger at the first abdominal segment and in concentric rings of the third leg. After *mlpt* RNAi *Ubx* is still strongly expressed in the first abdominal segment and not in concentric rings of the third leg. *Kr* RNAi embryos show diffuse staining in the abdominal region. Strong phenotype of *gt* RNAi embryos lack abdominal *Ubx* expression, while this marker is expressed in two consecutive pairs of legs, suggesting duplication of the third leg (L3). *Hb* RNAi embryos display large changes in the fate map with posterior abdominal expression of *Ubx* appearing at anterior (gnathal) segments. No *Ubx* staining was observed in the legs.

The expression of the Hox gene *Ultrabithorax* (*Ubx)* was also evaluated in control and in knockdowns of putative developmental genes. In *R. prolixus* control embryos, *Ubx* is strongly expressed in the first abdominal segment and in concentric rings around the third leg (L3) (Figure 11). As previously described, the effect of *mlpt* RNAi can generate at least three classes of embryonic phenotypes. Analysis of an embryo showing four pairs of legs (class 2 phenotype) showed that only the posterior-most leg expresses *Ubx*, suggesting that this leg corresponds to the L3 and that a duplication of L1 or L2 must have occurred. Interestingly, the expression of *Ubx* in the L3 does not occur as concentric ring as in the control, but rather in one side of the legs, suggesting that leg patterning was also affected (Figure 11). In addition, a slight expansion of *Ubx* towards posterior segments was observed in *mlpt* RNAi embryos (Figure 11). Immunohistochemical analysis using a monoclonal antibody against the homeobox transcription factor ANTP showed a staining in the thoracic segments and the leg one and two (L1 and L2) in the controls. In contrast, segmental fusion and segmental duplication with corresponding changes in ANTP staining were observed in *mlpt* RNAi embryos. Altogether, these results provided evidence that *mlpt* is locally important for the distinction between thoracic and abdominal identity.

*Ubx* expression was also analyzed upon knockdown of other gap genes such as *Kr*, *gt*, and *hunchback* (*Hb)* pRNAi. *Kr* RNAi embryos show only two pairs of legs and lack of *Ubx* staining in the legs, suggesting that the third leg (L3) is absent. In addition, a large expansion of *Ubx* expression towards the posterior region in *Kr* RNAi embryos was observed. *Rp-gt* RNAi embryos lack gnathal and abdominal segments and show a duplication of L3. *Rp-Hb* RNAi embryos show a large expansion of the posterior fates towards the anterior region, also pointing to a major role of this gap gene in *R. prolixus*. Altogether, these results suggest that the *Rp-mlpt* role in *R. prolixus* is rather limited and mainly concentrated at the transition between the thoracic and abdominal region and that it has a minor role in head morphogenesis, while gap genes such as *kr, gt, hb* display a broader role in the segmental cascade of the hemiptera *R. prolixus* (Figure 11).

## Discussion

### Nucleotide and peptide sequence analysis provides insights into the evolution of the mlpt gene function

In the past years, genes encoding smORFs have been characterized as new and important players of an unknown part of animal genomes [3; 8; 28]. *mlpt* is the most widely studied polycistronic gene encoding smORFs in insects [4; 6; 14; 15; 27; 28; 33]. Previous studies and our own sequence analysis presented here showed that *mlpt* is not present in chelicerate and myriapod genomes, being restricted to Pancrustacea.

Previous analyses identified specific roles for these smORF-encoded peptides (SEPs) peptides (SEPs) containing LDPTG motifs in the insect *D. melanogaster*, while a second weakly conserved predicted peptide (ORF-B in [5]) is not translated in *D. melanogaster* cell culture nor functionally required in rescue experiments [4; 5]. Mechanistically, small peptides from *mlpt* containing LDPTG motifs are essential for a selective proteasome-mediated N-terminal processing and activation of the transcription factor *svb* [27; 28].

Our docking and *in silico* mutation analysis identified the most important residues for the interaction of these small LDPTG peptides and the N-terminal region of the Svb protein, confirming previous experimental data in *D. melanogaster* [27; 28], and suggest a conserved interaction between small LDPTG peptides and Rp-Svb of *R. prolixus*.

The new hemipteran specific peptide identified here is approximately 35% longer in *R. prolixus* than in the most basally branching species analyzed, *Pseudococcus longispinus* (Figure 1 A, C). It has been recently proposed that smORFs could evolve to major ORFs via an “elongation” pattern [34]. Although smHemiptera is not a major ORF, size distribution of the smHemiptera peptide in hemiptera phylogeny corroborate this hypothesis. Interestingly, docking analyses of the larger hemiptera-specific predicted peptide (smHemiptera) were unable to detect specific molecular interactions between this peptide and the N-terminus of Svb, suggesting that this order-specific peptide might interact with other protein partners or other domains of Svb.

Importantly, the pRNAi technique used in our study leads to the knockdown of the mature transcript and presumably of all predicted peptides encoded by the hemipteran *mlpt* gene. Cas9/CRISPR editing technology has been established in other non-model arthropod species [35] and its establishment in *R. prolixus* might help to unveil the small peptides’ specific functions of *mlpt* in different developmental contexts.

Recent studies demonstrated that *mlpt* is required for Svb activation in adult tissues, particularly in nephric and intestinal *D. melanogaster* stem cells [36; 37]. Since *mlpt* is also expressed in *R. prolixus* digestive tract ([21] and our own observations), it will also be interesting to address the functional role of this gene containing smORFs in other developmental contexts.

### Conservation of mlpt expression among insects

*In situ* hybridization analysis of the *Rp-mlpt* gene during embryonic development shows a complex pattern of spatial expression, similarly to previously reported expression patterns in beetles, fruit flies and other holometabolous insects [5; 6; 13; 14; 33; 38]. Most differences in expression have been observed in early developmental stages. In *D. melanogaster*, the expression occurs in seven blastodermal stripes, a pair-rule pattern, while in most other insect species, including the basally branching Diptera midge *Clogmia albipunctata*, the expression appears as a gap domain [13]. Thus, as observed for *T. castaneum* [6] and more recently in the hemiptera *Oncopeltus fasciatus* [33], *Rp-mlpt* is first expressed in an anterior domain overlapping with the head and shortly after also in a posterior domain overlapping with the prospective thoracic segments (Figure 4). Later on, during germ band elongation and dorsal closure, the dynamic expression of *Rp-mlpt* is observed in several tissues including the tips of the legs, the head and the antenna (Figure 4). While expression in the head and in the legs have been reported in other species [5; 33], expression in the antennae at later stages has, to our knowledge, not been reported in other insect species.

Comparison of *Rp-mlpt* expression by RT-PCR between fertilized and non-fertilized eggs during the first hours of development (0-6 hours AEL) and 36-48 AEL shows ten times higher expression at later stages when compared to freshly laid eggs (0-6 hours AEL). In contrast, *Rp-svb* expression is higher during the initial stages of embryogenesis than in later stages, although its levels decrease by only 60% during these stages (Figure 5). Altogether, *Rp-mlpt* displays changes in extensive relative expression among the analyzed developmental stages and is spatially very dynamic, while the relative expression of *Rp-svb* is less variable among the developmental stages. These results suggest that *mlpt* levels are mediated by transcriptional control, and that the translated Mlpt peptides might post-transcriptionally switch Svb from a repressor to an activator at 36-48 hours AEL, as previously reported for *D. melanogaster* [27; 28].

Recent data suggest that an ancient regulatory system involving an Mlpt/Ubr3/Svb module operates in several insect species and overlapping expression domains of *mlpt* and *svb* have been reported [33]. Finally, *Rp-Notch* expression is downregulated upon *Rp-mlpt* RNAi (Figure 5). *In D. melanogaster, Notch* and *mlpt* have been shown to genetically interact during tarsal joint development. Therein, the Notch signalling pathway activates *mlpt* expression in each presumptive joint region. A feedback loop involving Svb and the Notch-ligand Delta is also involved in *mlpt* regulation [31; 32]. Thus far, studies on the role of Notch during *R. prolixus* segmentation are lacking and data on other insects is contradictory [32; 39; 40]. Future studies in *R. prolixus* should address the role of the Notch pathway in segmentation and its relationship with the transcription factor Svb.

### mlpt functions not as a classical gap gene in R. prolixus but rather participates in thoracic versus abdominal segmental identity

Three phenotypic classes were observed in *mlpt* knockdown embryos, mainly affecting segment formation (Figures 6 and 7). Comparison of the phenotypes of *mlpt* knockdown embryos with the phenotypes of transcription factor gap genes such as *Kr*, *Hb* and *Gt* (Figure 11) showed clear phenotypic differences. *mlpt* knockdown embryos display localized phenotypic changes mainly at the transition between thoracic and abdominal segments, while knockdown of the aforementioned transcription factors shows larger effects and changes in molecular marker expression. In *T. castaneum mlpt* knockdown leads to extensive homeotic transformations and embryos show up to six leg pairs [6]. Recent data in other hemiptera showed that knockdown of three genes *mlpt, Ubr3* and *svb* shows similar phenotypic effects as we describe here for *R. prolixus* such as posterior truncation, with the fusion/loss of thoracic segments, shortened legs and head appendages [33]. Phenotypes obtained by *Rp-svb* knockdown are similar to the *Rp-mlpt* RNAi embryos. Detailed analysis of the segment polarity gene *Rp-hh* showed that fusion takes place particularly at the thoracic regions and that abdominal segmentation and head formation is impaired in the strongest knockdown phenotypes (Figure 11). Analysis of the ventral midline gene *Rp-single-minded (Rp-sim)* showed that the ventral midline expression does not change upon *Rp-mlpt* knockdown, suggesting that dorsoventral patterning is not affected after *mlpt* RNAi (Sup. Figure 5).

Finally, several distal duplications of leg segments and leg malformations have been observed after *Rp-mlpt* RNAi (Figures 8 and 9). While several leg defects might be attributed to thoracic segment fusion, at least some examples of distal duplications are likely attributed to local effects of *Rp-mlpt* in the legs. We did not detect multiple rings of *Rp-mlpt* expression in *R. prolixus*; however, such rings were observed in *Periplaneta* but not in other hemipteran species [33; 41].

### Evolutionary crossroads at mlpt regulation

Recent studies revealed that *D. melanogaster mlpt* is regulated by ecdysone through direct binding of the nuclear ecdysone receptor (EcR) to its cis-regulatory region [15]. Transcription *mlpt* and translation of small peptides containing LDPTG motifs are then able to switch the role of Svb from a repressor to an activator. Our molecular docking data provides evidence suggesting that the interaction between the small peptides and Svb might be conserved in *R. prolixus.* Whether ecdysone plays a role during *R. prolixus* embryonic development is unknown, but data from another hemipteran species, the milkweed bug *Oncopeltus fasciatus*, suggests that at least one early ecdysone-responsive gene, the transcription factor *E75* is expressed as a pair-rule gene and generates similar phenotypes to the ones described here for *mlpt* with thoracic segmental fusion and lack of abdominal segments [42]. As described for the role of *mlpt* in segmentation ([33] and here)*, E75* is also involved in the same process in *O. fasciatus*, while its role in segmentation was lost in the lineage giving rise to the Diptera *D. melanogaster* (Figure 12). It is not clear if ecdysteroid titers change during *R. prolixus* embryogenesis and if ecdysone directly regulates *mlpt* expression during these stages. Interestingly, after *Rp-mlpt* parental RNAi a few nymphs hatch and show defective leg hardening and darkening, suggesting that cuticle has been affected. Previous studies [42] and our data presented here suggest possible connections between the segmentation and molting gene networks, which should be addressed by future studies.

**Figure 12:**
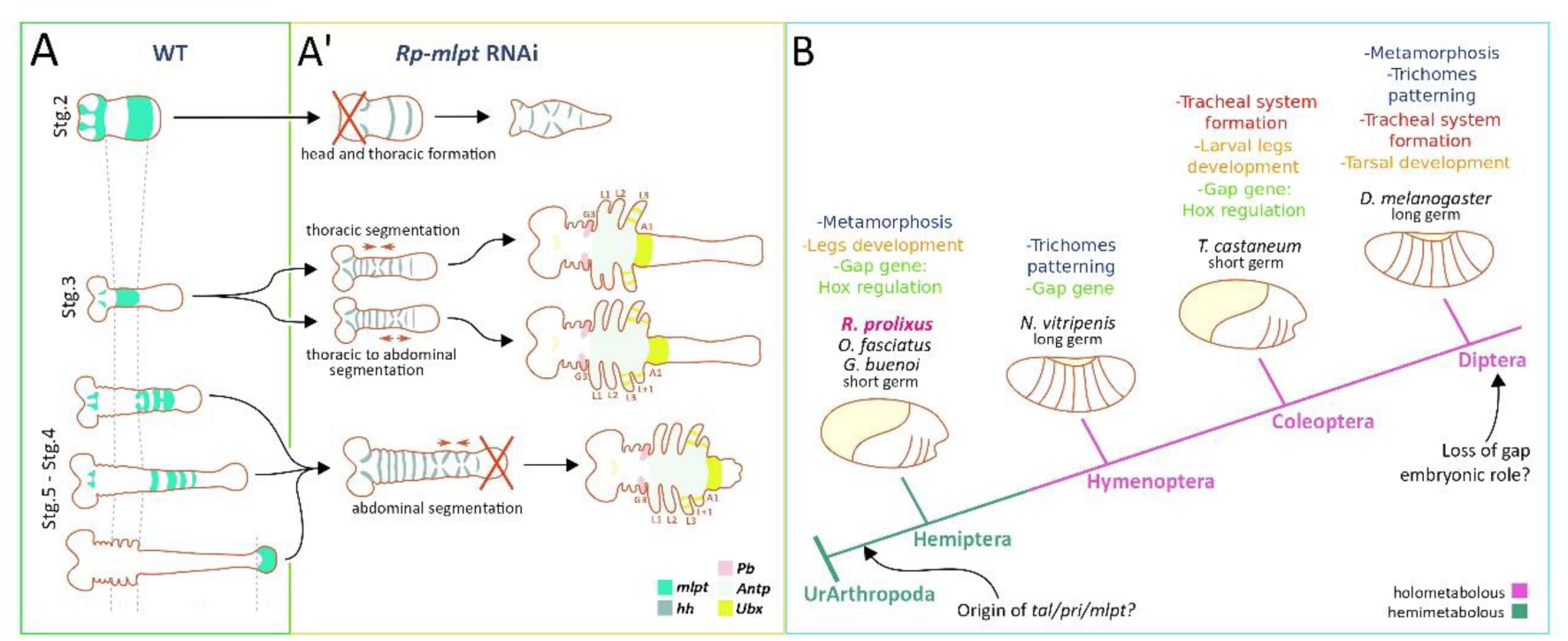
Model of *mlpt* roles during *R. prolixus* embryonic development. (A) *mlpt* is required for the correct specification of the thoracic segments and patterning of the distal structures of the legs and in the maintenance of the head regions. (B) Multiple roles of *mlpt* during insect evolution based on findings of [33] and the current manuscript.

## Material & Methods

### Bioinformatic analyses

Mlpt peptide sequences from *D. melanogaster* and *T. castaneum* were used for BLAST searches [43] against available arthropod genomes and transcriptomes using relaxed parameters to maximize the chances to obtain genes encoding smORFs. The species used for the smORF identification are depicted in Sup. Table 1. *Rp-mlpt/tal/pri* was identified in *R. prolixus* genome and transcriptomes [19; 20; 21; 22]. BLAST results and domain architectures were manually annotated. For regulatory sequence *in silico* searches in the *D. melanogaster, T. castaneum* and *R. prolixus* genome a strategy similar to [44] was designed. The *D. melanogaster* Ecdysone Receptor motif was obtained in the FlyFactorSurvey database [45] and used to scan upstream, intronic and downstream of *mlpt* genomic regions of *D. melanogaster, T. castaneum* and *R. prolixus* using FIMO [46].

### Structural modeling and Shavenbaby-MLPTs docking assays

The 3D models of Dmel-Svb (834 aa; # CAD23206.1) and Rprol-Svb (834 aa; # mRNA: GECK01059228) were predicted via *ab initio* modeling using I-Tasser program [47]. The 3D structures of MLPTs (Dmel-pptd1 and 4) were predicted via CABS-dock and I-Tasser [48], respectively; Rprol-pptd1 and 2, both via CABS-dock. The global and local stereochemical quality of all predicted models were performed via Rampage [49], Verify3D [50], ProSA [51], VoroMQA [52], ProQ3D [53], Qprob [14], DeepQA [54] and SVMQA [55]. The most suited structural models were refined via ModRefiner [56]. The protein-peptide docking assays were directed to the Shavenbaby N-terminal region (31 aa initial) inspired on the findings of [28]. The dockings were performed using ClusProV2 [57], HADDOCK [58], SwarmDock [59] and HDock [60] programs. The best interaction complexes were selected after exhausting analyzes with DockScore [61], PPCheck [62] and CCharPPI [63] programs. The best solutions for Dmel-Svb-pptd1 and 4 complexes were obtained via Haddock and ClusProV2, respectively; for Rprol-Svb-pptd1 and 2 complexes, via SwarmDock and Haddock, respectively. After selection of the best complex, further refinements were performed via GalaxyRefineComplex [64]. The refinements were completed using the “*Energy minimization of side chains*” function on PDB_Hydro webserver [65]. The prediction of free energy (ΔG), dissociation constant (*K*_d_) as well as the analysis of hotspot amino acids located in the interaction interface, were performed via PRODIGY [58; 66], HotRegion [67] and ANCHOR [68], respectively. Calculations on the change in the binding energy of the protein-protein complex following mutations in the residues on interaction interface were performed via BindProfX [69]. Interatomic contacts and types of interactions maintained at Svbs-MLTPs interfaces were analyzed via PDBsum [70]. The images of Svbs-MLTPs complexes were obtained via UCSF Chimera 1.13.1 [71].

### Insect rearing, fixation and dissection

Insect rearing was performed as described by [72]. Approximately one week after blood-feeding eggs were collected daily and fixed in different stages of development. For fixation, up to 100 eggs were briefly washed with distilled water to remove the debris and then transferred to a 1.5 mL microtube containing 1 mL of distilled water. This microtube was maintained for 90 seconds in boiling water, and after this period the water was replaced by 1 mL of paraformaldehyde 12% (PBS) and fixed for one hour (6-8°C). Then, the embryos were incubated with 1 mL of paraformaldehyde 4% containing 0,1% of Tween 20 under agitation (200rpm) for one hour at room temperature. The eggs were then washed repeatedly with PBST (PBS 1X, Tween 20 0.1%). For long-term storage, embryos were gradually transferred to ethanol 100% and then stored at −20°C. For the dissections two forceps (Dumont No. 5) were used. The egg is hold with the help of one of them, while the other is used to pressure at the chorionic rim to remove the operculum. The chorionic rim is hold and the shell is opened transversely, leading to the embryonic release. For later stages, when segmentation is complete, at least stage 6 [23], the yolk is easily removed either with the help of a thin forceps or a glass needle, without damaging the embryo.

### In situ hybridization, nuclear and antibody staining

Embryos stored in ethanol 100% at −20°C were gradually transferred to PBST at room temperature. *in situ* hybridization and probe synthesis were performed as described by [73] including the proteinase K treatment. DAPI (4′,6-Diamidine-2′-phenylindole dihydrochloride, SIGMA) staining was performed as in [23]. All the images were acquired with the stereoscope Leica M205 and processed and analyzed with the software Leica Application Suite Advanced Fluorescence Version 0.4 (LAS AF v4 - Leica Microsystems). Images were assembled and the *in situ* hybridization staining (NBT/BCIP) was converted to a false fluorescence as described in [74].

### RNA interference and real-time PCR

The RNA interference (RNAi) was performed similarly to [44] using a non-related dsRNA as a control (neomicin dsRNA-dsneo). For *mlpt*, two non-overlapping PCR fragments containing T7 promoter initiation sites at both ends were used as templates for dsRNA synthesis using Ambion T7 Megascript Kit (Cat. No. AM1334). The amount and integrity of the dsRNA samples were measured by spectrophotometry and agarose gel electrophoresis, respectively. For each silenced gene between 6 to 12 micrograms of dsRNA were injected. The quantification of RNAi efficiency and comparison of gene expression after silencing was measured through real-time PCR (RT-PCR) as performed in [23] using the gene Elongation factor-1 (Ef1) as endogenous reference gene. Gene bank or Vector base accession numbers of the genes analyzed by *in situ* hybridization or RT-PCR are provided in the Supplementary table 3. For RT-PCR analysis, RNA was extracted in biological triplicates from eggs at appropriate hours after egg lay (AEL). Fertilized eggs (0-6 hours and 36-48 hours AEL) and non-fertilized eggs obtained from female virgins (0-6 hours AEL) were used for the analysis.

## Acknowledgements

RNdF is supported by CNPq (307952/2017-7 and 431354/2016-2) and FAPERJ (E-26/210-150/2016 and E-26/203.298/2016). VTdS and DGA were master students of PPG-PRODBIO-Macaé (CAPES scholarships), MB a PhD student from PPG-PCM-ICB-UFRJ and LR was a postdoc with CAPES scholarship from the National Institute of Molecular Entomology/CNPq (INCT-EM). The authors thank Roland Zimm for helpful suggestions in the manuscript.

