## Supplementary File for "Multiple roles of the polycistronic gene *tarsaless/mille-pattes/polished-rice* during embryogenesis of the kissing bug *Rhodnius prolixus*"

**Running title:** *tal/mlpt/pri in the hemiptera Rhodnius*

| Species | Family | Order | Genbank Accession Number | Transcript Length (bp) |
|---|---|---|---|---|
| *Rhodnius prolixus* | Reduviidae | Hemiptera | GECK01030379 | 1,323 |
| *Triatoma pallidipennis* | | | GEVD01005550 | 953 |
| *Cimex lectularius* | Cimicidae | | GBYH01024474 | 1,666 |
| *Lygus hesperus* | Miridae | | GBHO01010581 | 1,509 |
| *Chinavia ubica* | Pentatomidae | | GBFA01008810 | 550 |
| *Clavigralla tomentosicollis* | Coreidae | | GAJX01001486 | 1,331 |
| *Pyrrhocoris apterus* | Pyrrhocoridae | | GFOX01081802 | 751 |
| *Riptortus pedestris* | Alydidae | | AK417152 | 1,577 |
| *Acanthosoma haemorrhoidale* | Acanthosomatidae | | GAUV02015318 | 616 |
| *Velia caprai* | Veliidae | | GAUO02023323 | 819 |
| *Graminella nigrifrons* | Cicadellidae | | GAQX01032246 | 1,386 |
| *Cercopis vulnerata* | Cercopidae | | GAUN02002429 | 866 |
| *Okanagana villosa* | Cicadidae | | GAWQ02039649 | 982 |
| *Essigella californica* | Lachnidae | | GAZF02055449 | 2,318 |
| *Acyrthosiphon pisum* | Aphididae | | NM_001134487 | 1,751 |
| *Pseudococcus longispinus* | Pseudococcidae | | FIZU01058922 | 51,350 (contig) |
| *Tribolium castaneum* | Tenebrionidae | Coleoptera | NM_001114384 | 616 |
| *Aedes aegypti* | Culicidae | Diptera | GFNA01046000 | 2,370 |
| *Drosophila melanogaster* | Drosophilidae | | JV223590 | 917 |
| *Timema cristinae* | Timematidae | Phasmatodea | GAVX02010179 | 855 |
| *Epiophlebia superstes* | Epiophlebiidae | Odonata | GAVW02017495 | 838 |
| *Folsomia candida* | Isotomidae | Entomobryomorpha | GASX02021174 | 373 |
| *Bragasellus molinai* | Asellidae | Isopoda | HAEM01125628 | 474 |

**Supplementary Table 1: Species, family and order names, accession numbers and transcript lengths (base pairs) of *mlpt* related transcripts.**

| Organism | Protein | No. of interface residues | Interface area (Å$^2$) | Hotspot | No. of salt bridges | No. of hydrogen bonds | No. of non-bonded contacts | Binding affinity (ΔG) | Dissociation Constant (Kd) (M) |
|---|---|---|---|---|---|---|---|---|---|
| Dmel | Svb | 23 | 678 | F5, I7, K23 | - | 4 | 94 | -10.9 | 1.1E-08 |
| | pptd1 | 10 | 845 | L5, D6, Y11 | | | | | |
| Dmel | Svb | 23 | 1050 | M1, P2, E26 | 5 | 7 | 127 | -11.5 | 3.9E-09 |
| | pptd4 | 18 | 1212 | T7, R10, R12, R21 | | | | | |
| Rprol | Svb | 17 | 649 | L11, Q13, Q15, L16 | - | 5 | 81 | -9.3 | 1.5E-07 |
| | pptd1 | 10 | 866 | N4, L6, L11, Y12 | | | | | |
| Rprol | Svb | 26 | 864 | K3, L12, Q15 | - | 6 | 133 | -10.0 | 4.7E-08 |
| | pptd2 | 15 | 1041 | M2, P11, Y15 | | | | | |

**Supplementary Table 2: Molecular docking interaction profile between Svb and Mlpt peptides. Dmel - *Drosophila melanogaster,* Rprol - *Rhodnius prolixus*.**

| Gene name | Acession number: | Forward Primer | Reverse Primer | Product size |
|---|---|---|---|---|
| *Rp-mille-pattes* | GECK01030379.1 (reverse complement) | ggccgcggTTTGGACCC AACAGGTCTTT | cccggggcGGTGTCCCCTA TTGGTCCTT | 441 bp |
| *Rp-hedgehog* | VB: RPRC012384-RA | ggccgcggGTAGATCTG AGGCGGATTGC | cccggggcCAGAAATTGGG TGCGCTACT | 631 bp |
| *Rp-Krüppel* | VB:RPRC000102-RA | ggccgcggCTATGGCTA GGCGAGAACCA | cccggggcCTGTTCAGGGA GGTCGAGAG | 791 bp |
| *Rp-giant* | VB: ACPB03018756.1 | ggccgcggTACACCACA GGCTTCACAGG | cccggggcAAAAGCCGCTC GTATAGCAA | 555 bp |
| *Rp-hunchback* | VB: ACPB03005501.1 | ggccgcggCATGCCCAA AGTGTCCTTTT | cccggggcAAGCATCCGTG CTGTTCTCT | 600 bp |
| *Rp-shavenbaby* | RPRC002781-RA | ggccgcggTCAGATCTT GATGGCTGCTG | cccggggcTTGCCATTCAC TCTGTGCTC | 813 bp |
| *Rp-proboscipedia* | FJ694871.1 | ggccgcggTTGGCCTCA TGTTATCAGCA | cccggggcGTAGATCTGAG GCGGATTGC | 666 bp |
| *Rp-Ultrabithorax* | RPRC000565-RA | ggccgcggTCGAACAG ACGGGCTTTTAC | cccggggcACAAAGCGTGA GCCATTTCT | 762 bp |
| *Rp*-single-*minded* | ACPB03006262.1 | ggccgcggACCAGAAT GTCGGCCTAGTG | cccggggcCGGTTGATGAT GGTGATGAG | 845bp |
| *Rp-shavenbaby Real-time PCR* | RPRC002781-RA | AATTCCACCACCTGT TGAGC | TGCTTTGGCTGTGGTAC TTG | 213 bp |
| *Rp-mille-pattes Real-time PCR* | GECK01030379.1 (reverse complement) | TTTAGCTCGGACTCCT TTGG | GGTAGGATCGAGCGTA TTGG | 150 bp |
| *Rp-Notch* | Previously published in [1] | Previously published in [1] | Previously published in [1] | Previously published in [1] |

**Supplementary Table 3: Gene names, accession numbers and primer sequences of *R. prolixus* selected sequences with respective product sizes.** Lowercase letters in the primers correspond to adaptor sequences for sense (ggccgcgg) and anti-sense (cccggggc) T7 polymerase promoter sequences, as previously described [2; 3].

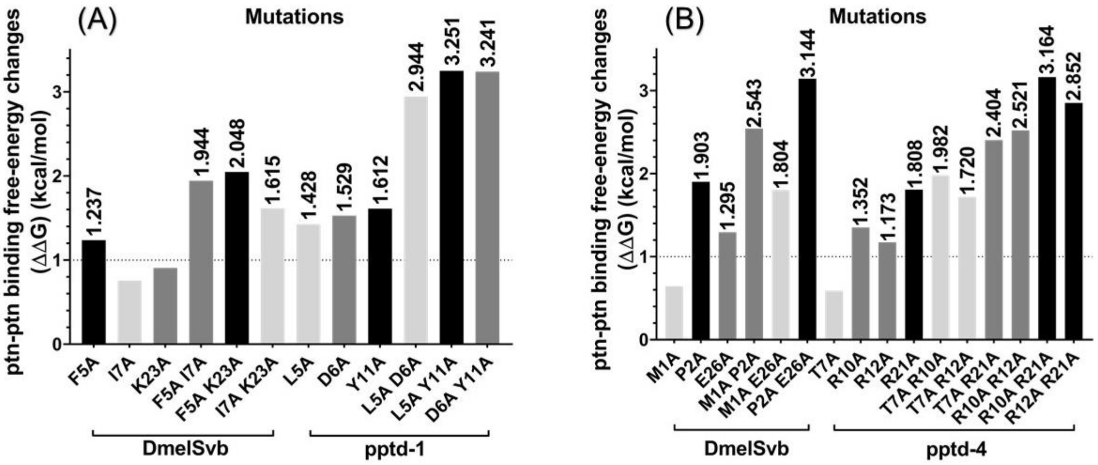


**Supplementary Figure 1: In silico single- and double-mutations on key amino acids at the Svb-peptide interface change the protein-peptide complex binding affinity.** The BindProfX tool was used to predict the protein-protein binding affinity change upon mutation of residues at the interface between the transcription factor DmelSvb and the small peptide Dmel-pptd1(A) and Dmel-pptd4 (B). (A) Largest effects (higher free-energy changes) were observed upon double mutations on the most conserved residues of Dmel-pptd1 (L5, D6, and Y11) and Dmel-pptd4 (R10, R12, and R21). Amino acids are represented in universal single-letter code. Dmel - *Drosophila melanogaster*.

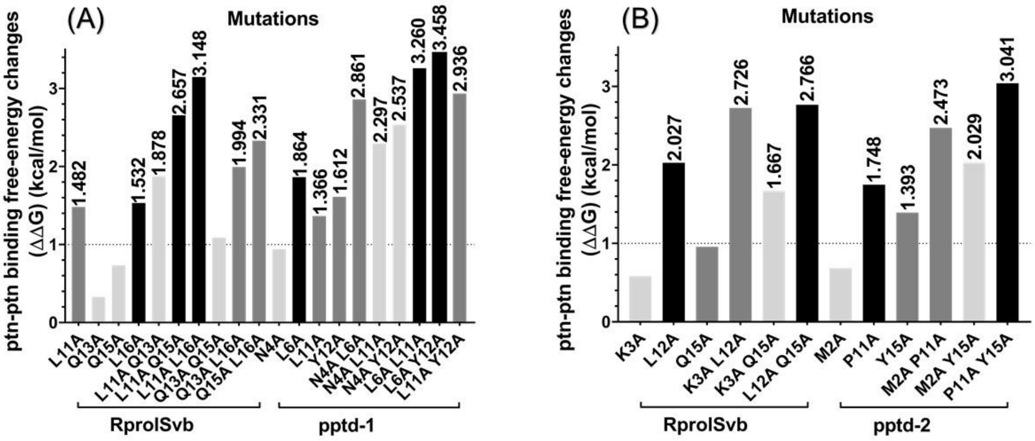


**Supplementary Figure 2: In silico single- and double-mutations on key amino acids at the Svb-peptide interface change the protein-peptide complex binding affinity.** The BindProfX tool was used to predict the protein-protein binding affinity change upon mutation of residues at the interface between the transcription factor RprolSvb and the small peptide Rprol-pptd1(A) and Rprol-pptd2 (B). (A) Largest effects (higher free-energy changes) were observed upon double mutations on most conserved residues of Rprol-pptd1 (N4, L6, L11 and Y12) and Rprol-pptd2 (M2, P11 and Y15). Amino acids are represented in universal single-letter code. Rprol - *Rhodnius prolixus*.

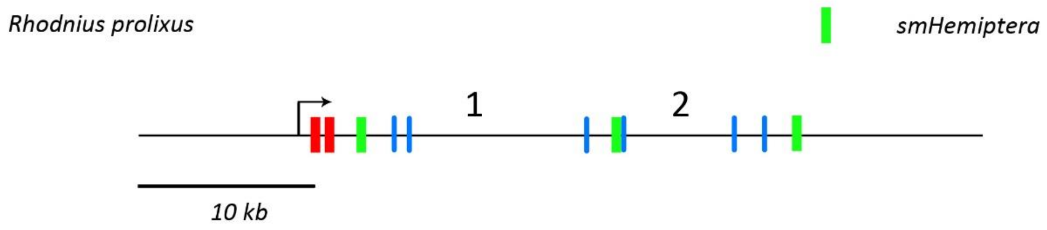


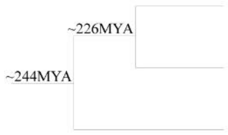


| | Intron 1 | Intron 2 | CDS 1 | CDS 2 | CDS 3 |
|---|---|---|---|---|---|
| ***Rhodnius prolixus*** | 17.185 nt | 10.418 nt | 57 nt | 123 nt | 69 nt |
| ***Cimex lectularius*** | 1.664 nt | 376 nt | 57 nt | 111 nt | 66 nt |
| ***Halyomorpha halys*** | 29.491 nt | 6.425 nt | 45 nt | 96 nt | 69 nt |
| **Identity** | ~0% | ~0% | 33.3% | 23.2% | 61.1% |

**Supplementary Figure 3: Intron and CDS size of smHemiptera in three Hemipteran species.** *Rhodnius prolixus* locus is constituted by *tal/mlpt* CDS (red boxes) and smHemiptera CDS (green boxes). 1 and 2 refers to introns 1 and 2 of smHemiptera. CDS identity is higher than intronic conservation in all species. CDS size is similar among the three analyzed species, while intron size appears more variable.

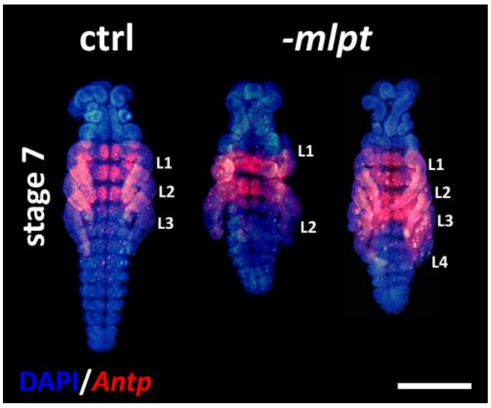


**Supplementary Figure 4: ANTP antibody staining in control and in *Rp-mlpt* RNAi embryos**. ANTP antibody marks the first and the second legs (L1 and L2) and the three thoracic segments in control embryos. Middle row - *Rp-mlpt* embryos show two fused segments (L1 and L2) and a third leg (L3), which does not express ANTP. Rightmost embryo shows four legs with a probable duplication of L3, since the two most posterior legs do not express ANTP.

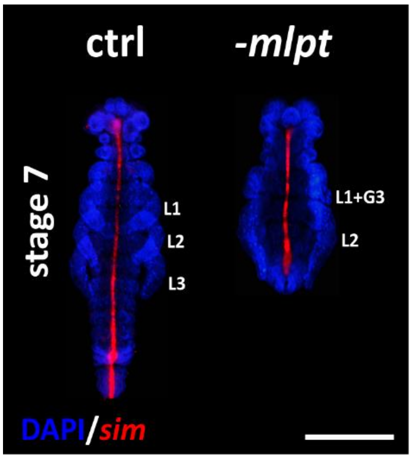


**Supplementary Figure 5: *Rp-sim* expression in control and *Rp-mlpt* RNAi embryos.** *Rp-single-minded* is expressed in the ventral midline of control and *Rp-mlpt* embryos. No significant difference in expression of *Rp-sim* was observed in both embryonic groups.

[1] F.B. Mury, M.D. Lugon, D.A.F. RN, J.R. Silva, M. Berni, H.M. Araujo, M.R. Fontenele, L.A. Abreu, M. Dansa, G. Braz, H. Masuda, and C. Logullo, Glycogen Synthase Kinase-3 is involved in glycogen metabolism control and embryogenesis of Rhodnius prolixus. Parasitology 143 (2016) 1569-79.

[2] L. Ribeiro, V. Tobias-Santos, D. Santos, F. Antunes, G. Feltran, J. de Souza Menezes, L. Aravind, T.M. Venancio, and R. Nunes da Fonseca, Evolution and multiple roles of the Pancrustacea specific transcription factor zelda in insects. PLoS Genet 13 (2017) e1006868.

[3] M. Berni, M.R. Fontenele, V. Tobias-Santos, A. Caceres-Rodrigues, F.B. Mury, R. Vionette-do-Amaral, H. Masuda, M. Sorgine, R.N. da Fonseca, and H. Araujo, Toll signals regulate dorsal-ventral patterning and anterior-posterior placement of the embryo in the hemipteran Rhodnius prolixus. Evodevo 5 (2014) 38.
